# Cancer cell CCR2 orchestrates suppression of the adaptive immune response

**DOI:** 10.1101/390187

**Authors:** Miriam R. Fein, Xue-Yan He, Ana S. Almeida, Emilis Bružas, Arnaud Pommier, Ran Yan, Anaïs Eberhardt, Douglas T. Fearon, Linda Van Aelst, John Erby Wilkinson, Camila O. dos Santos, Mikala Egeblad

## Abstract

C-C chemokine receptor type 2 (CCR2) is expressed on monocytes and facilitates their recruitment to tumors. Though breast cancer cells also express CCR2, its functions in these cells are unclear. We found that *Ccr2* deletion in cancer cells led to reduced tumor growth and ∼2-fold longer survival in an orthotopic, isograft breast cancer mouse model. Deletion of *Ccr2* in cancer cells resulted in multiple alterations associated with better immune control: increased infiltration and activation of cytotoxic T lymphocytes (CTLs) and CD103+ cross-presenting dendritic cells (DCs), as well as upregulation of MHC class I and downregulation of checkpoint regulator PD-L1 on the cancer cells. Pharmacological or genetic targeting of CCR2 increased cancer cell sensitivity to CTLs and enabled the cancer cells to induce DC maturation toward the CD103+ subtype. Consistently, *Ccr2^−/−^* cancer cells did not induce immune suppression in *Batf3^−/−^* mice lacking CD103+ DCs. Our results establish that CCR2 signaling in cancer cells can orchestrate suppression of the immune response.

**Summary:** C-C chemokine receptor type 2 (CCR2) expressed on monocytes facilitates their recruitment to tumors. Here, CCR2 signaling in cancer cells is shown to suppress immune control of tumors, in part by reducing CD103+ dendritic cell recruitment.

## INTRODUCTION

Tumors escape immune control via multiple mechanisms (Dunn et al., 2002; Dunn et al., 2004). These mechanisms include cancer cell-intrinsic changes that alter how the cancer cell is recognized by the immune system and extrinsic changes that suppress immune cell activities. For example, cancer cells can intrinsically decrease the surface expression of major histocompatibility complex (MHC) class I, making them effectively invisible to cytotoxic T lymphocytes (CTLs) (Garrido et al., 2016). Moreover, cancer cells can upregulate programmed cell death ligand 1 (PD-L1/B7-H1), which binds the PD-1 receptor on activated T cells, resulting in T cell anergy, ultimately protecting cancer cells against T cell-mediated killing (Chen et al., 2016a). Extrinsic mechanisms include the downregulation of co-stimulatory molecules (*e.g.*, CD86) on antigen-presenting cells; the secretion of cytokines that directly inhibit CTLs; and the recruitment of regulatory T cells and myeloid-derived suppressor cells (MDSCs) (Igney and Krammer, 2002). In contrast, infiltration of CD103+ dendritic cells (DCs) in mice, phenotypically similar to CD141+ DCs in humans (Hildner et al., 2008), has emerged as a mechanism by which tumors may be kept under immune control. CD103+ DCs are highly efficient at acquiring and processing exogenous antigens, which they directly present on MHC class I molecules to CD8+ CTLs. Even a modest accumulation of CD103+ DCs in tumors has been associated with improved immune-mediated tumor control (Broz et al., 2014; Roberts et al., 2016).

Chemokine receptors mediate the recruitment of immune cells to sites of inflammation and to tumors. C-C chemokine receptor type 2 (CCR2) is expressed by several bone marrow-derived cell types (including inflammatory monocytes, myeloid precursor cells, and immature DCs), as well as B and T lymphocytes (Lim et al., 2016). CCR2-expressing cells are recruited to sites of inflammation primarily by C-C chemokine ligand 2 (CCL2) (Deshmane et al., 2009), although other CCL family members also can activate CCR2. Chemotherapy treatment promotes CCR2-dependent infiltration of tumor-promoting myeloid cells to murine mammary tumors (Nakasone et al., 2012), and CCL2/CCR2-mediated recruitment of CCR2+ inflammatory monocytes to the lung has been shown to promote breast cancer extravasation and metastasis in mouse models (Qian et al., 2011). Furthermore, elevated levels of CCL2 in tumors and in serum are associated with advanced disease and poor prognosis in breast carcinoma patients (Lebrecht et al., 2004; Lebrecht et al., 2001; Soria and Ben-Baruch, 2008). These findings have sparked interest in targeting the CCR2 pathway to modulate the innate immune response for therapeutic benefit in cancer. However, CCR2 is also expressed by breast cancer cells, and activation of CCR2 by CCL2 can induce cancer cell migration and survival through Smad3-, p42/44MAPK-, and Rho GTPase-mediated signaling (Fang et al., 2012). *In vivo*, the potential roles of CCR2 signaling in cancer cells have not been well investigated, largely because they were thought to be minor compared to the roles of CCR2 in myeloid cells.

In this study, we report that CCR2 signaling in cancer cells plays a surprisingly major role in regulating the immune response to murine breast tumors. We show that CCR2 in cancer cells supports immune escape by inhibiting CD103+ DC infiltration and maturation and by suppressing cytotoxic T cell activity. CCR2 expression on cancer cells represents a previously uncharacterized mechanism for immune suppression. Thus, our data support the notion that the CCL2/CCR2 axis is an important immune modulatory pathway in cancer, utilized by both immune cells and cancer cells to orchestrate immune suppression.

## RESULTS

### CCR2 in cancer cells promotes primary tumor growth in an orthotopic, isograft breast cancer mouse model

To investigate the potential effects of CCR2 on breast tumor growth and metastasis, we crossed *Ccr2*^−/−^ mice (Boring et al., 1997) with mouse mammary tumor virus - polyoma middle T (MMTV-PyMT) mice, a model of luminal B breast cancer (Guy et al., 1992; Lin et al., 2003). The *Ccr2* genotype did not influence normal mammary gland development (**Fig. S1a**) or tumor onset (**Fig. 1a**); however, loss of even one allele of *Ccr2* significantly reduced tumor growth rates (**Fig. 1b**). Consistently, survival time was significantly longer for the MMTV-PyMT;*Ccr2^−/−^* and MMTV-PyMT;*Ccr2^+/−^* mice compared to that of the MMTV-PyMT;*Ccr2^+/+^* mice (**Fig. 1c**). In addition, we noted that the tumors of MMTV-PyMT;*Ccr2^−/−^* and MMTV-PyMT;*Ccr2^+/−^* mice were more cystic, with reduced solid areas, than those of MMTV-PyMT;*Ccr2^+/+^* mice (**Fig. S1b, c**).

**Fig. 1:**
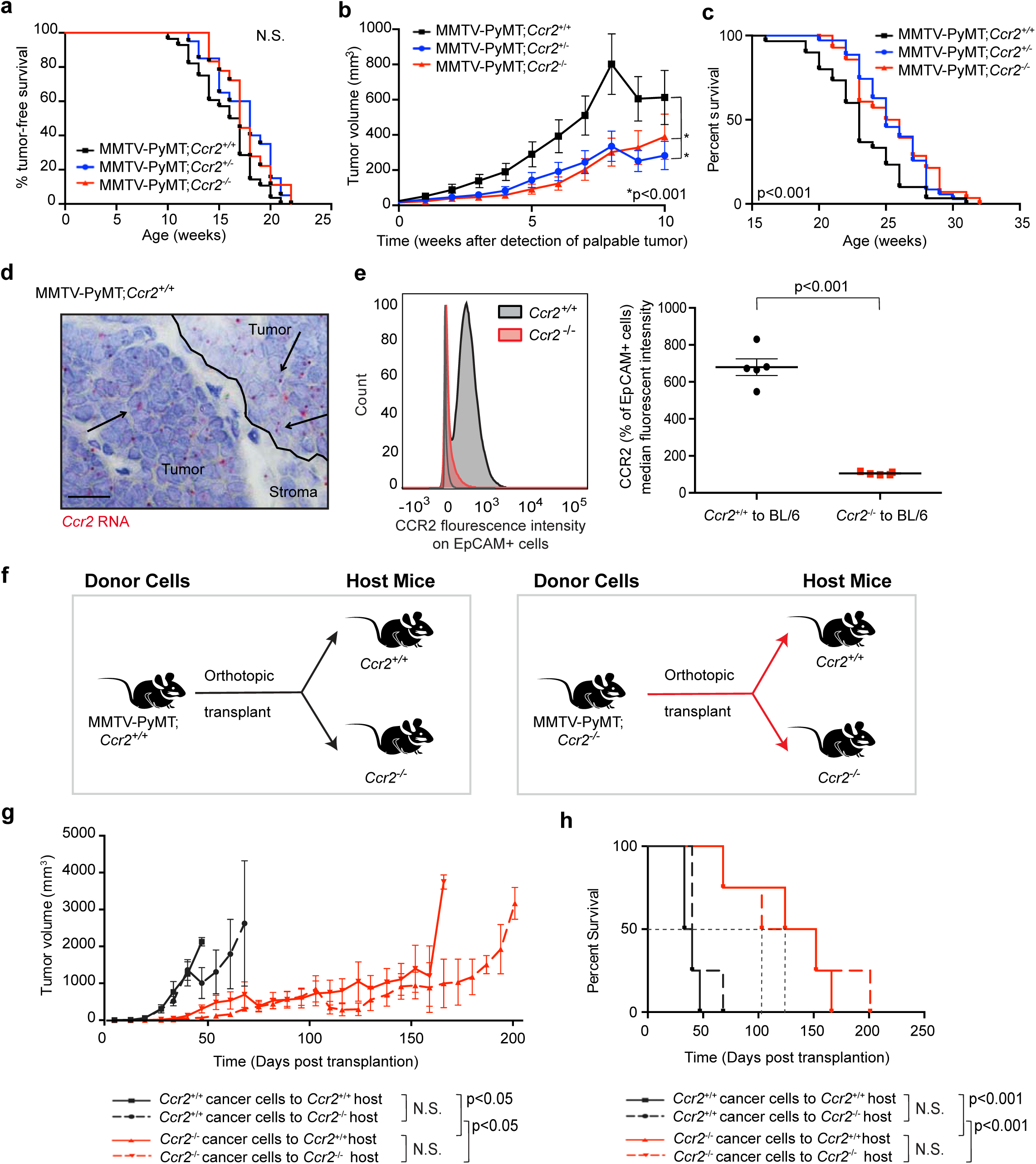
Cancer cell *Ccr2* promotes primary tumor growth, reducing overall survival. **a.** Tumor onset (tumor-free survival) is similar for MMTV-PyMT;*Ccr2*^+/+^, MMTV-PyMT;*Ccr2*^+/−^, and MMTV-PyMT;*Ccr2*^−/−^ mice, as determined by weekly palpation and caliper measurement. N.S.=non-significant (Log-rank [Mantel-Cox] test; n=28 MMTV-PyMT;*Ccr2*^+/+^, 20 MMTV-PyMT;*Ccr2*^+/−^, and 18 MMTV-PyMT;*Ccr2*^−/−^ mice). **b.** Primary tumor growth is reduced in MMTV-PyMT;*Ccr2*^+/−^ and MMTV-PyMT;*Ccr2*^−/−^ mice compared to MMTV-PyMT;*Ccr2*^+/+^ mice, as determined by weekly caliper measurement (mean +/− SEM, two-way ANOVA; n=21 MMTV-PyMT;*Ccr2*^+/+^, 25 MMTV-PyMT;*Ccr2*^+/−^, and 18 MMTV-PyMT;*Ccr2*^−/−^ mice). **c.** Time until IACUC-approved endpoint was increased for MMTV-PyMT;*Ccr2*^+/−^ and MMTV-PyMT;*Ccr2*^−/−^ mice compared to MMTV-PyMT;*Ccr2*^+/+^ mice (Log-rank [Mantel-Cox] test; n=30 MMTV-PyMT;*Ccr2*^+/+^, 35 MMTV-PyMT;*Ccr2*^+/−^, and 29 MMTV-PyMT;*Ccr2*^−/−^ mice). **d.** Cancer cells express CCR2 (arrows) as determined by RNA *in situ* hybridization for *Ccr2* performed on paraffin-embedded tumor sections from MMTV-PyMT;*Ccr2*^+/+^ mice. Image is representative of more than 10 mice. Scale bar=20 μm. **e.** CCR2 is expressed on epithelial cells, as determined by flow cytometry performed on *Ccr2^+/+^* or *Ccr2^−/−^* tumors transplanted to C57BL/6 hosts. Anti-CCR2 primary antibody was gated on EpCAM+ cells and compared to background levels. Left panel shows one representative experiment and right panel depicts median fluorescence intensity for all experiments (mean +/− SEM, Student’s t-test; n=5). **f.** Schematic of transplantation experiments. **g.** Primary tumor growth of *Ccr2*^−/−^ cancer cells is reduced compared to that of *Ccr2*^+/+^ cancer cells regardless of the *Ccr2* genotype of the host, as determined by weekly caliper measurement (mean +/− SEM, two-way ANOVA; n=8 for all conditions). **h.** Survival until IACUC-defined endpoint is increased in mice bearing *Ccr2*^−/−^ cancer cells regardless of the genotype of the host (Log-rank [Mantel-Cox] test; n=8 for all conditions).

The importance of CCR2 signaling for monocyte recruitment is well understood, but CCR2 is also expressed by human breast cancer cells (Fang et al., 2012). In accordance with this report, we detected *Ccr2* mRNA and CCR2 protein in the cancer cells of the tumors developing in MMTV-PyMT;*Ccr2^+/+^* mice (**Fig. 1d, e**), and using RNA fluorescence *in situ* hybridization (FISH), we demonstrated that *Ccr2* mRNA is expressed by *Krt18*-positive breast cancer cells (**Fig. S1d**). To determine the relative contributions of cancer cell vs. host CCR2 to tumor growth, we isolated primary cancer cells (purity >90%; **Fig. S1e**) from MMTV-PyMT;*Ccr2*^+/+^ and MMTV-PyMT;*Ccr2*^−/−^ mice and transplanted them orthotopically to mammary glands of either *Ccr2*^+/+^ or *Ccr2^−/−^* syngeneic host mice (**Fig. 1f**). Loss of *Ccr2* in the host did not alter tumor growth, consistent with our previous report (Nakasone et al., 2012). However, loss of *Ccr2* in cancer cells significantly reduced tumor growth rates, leading to ∼2-fold longer survival (**Fig. 1g, h**). *Ccr2* mRNA levels of *Ccr2^+/−^* cancer cells were lower than those of *Ccr2^+/+^* cancer cells (**Fig. S1f**), and tumors derived from *Ccr2^+/−^* cancer cells exhibited the same slow growth as tumors derived from *Ccr2^−/−^* cancer cells (**Fig. S1g**). This result suggests that a threshold level of CCR2 expression in MMTV-PyMT cancer cells regulates tumor growth.

CCR2+ inflammatory monocytes can promote breast cancer metastasis (Muller et al., 2001; Qian et al., 2011). The overall metastatic burden in the lung—the primary site of metastasis in the MMTV-PyMT model—was, however, very variable in MMTV-PyMT;*Ccr2^+/+^* mice and was not significantly different from that of MMTV-PyMT;*Ccr2^−/−^* mice (**Fig. S1h**). There were also no significant differences in the number of metastatic foci (representing seeding density) related to the *Ccr2* genotype (**Fig. S1i**). However, we found that the metastatic foci were larger in the lungs of MMTV-PyMT;*Ccr2^+/+^* mice than in the lungs of MMTV-PyMT;*Ccr2^−/−^* mice (**Fig. S1j**), suggesting that CCR2 promotes the growth of secondary lesions.

### *Ccr2* expression in cancer cells is associated with poor differentiation and reduced apoptosis sensitivity

The tumors from transplanted *Ccr2*^+/+^ or *Ccr2^−/−^* cancer cells were markedly different at the histological level (**Fig. 2a**), similar to the autochthonous tumors in the MMTV-PyMT;*Ccr2^−/−^* animals (**Fig. S1b, c**). The tumors originating from *Ccr2*^+/+^ cancer cells consisted entirely of microlobules composed of sheets of large, neoplastic epithelial cells. The microlobules were surrounded by small amounts of fibrovascular stroma containing dense collagen, which is characteristic of poorly differentiated MMTV-PyMT tumors. The neoplastic cells had round to oval nuclei, prominent nucleoli, scant cytoplasm, and indistinct cell borders, typical of undifferentiated cells. In contrast, the tumors originating from *Ccr2^−/−^* cancer cells were more differentiated. In many areas, the cancer cells contained lipid vacuoles of various sizes. These tumors also had large cystic areas lined by a single or double layer of small, polarized epithelial cells, and the lumina were filled with proteinaceous secretions (**Fig. 2a**). Conversely, the tumors originating from *Ccr2^−/−^* cancer cells had a reduced percentage of solid areas compared to those originating from *Ccr2^+/+^* cancer cells (**Fig. 2b**).

To understand why tumors from *Ccr2^−/−^* cancer cells grew slower than tumors from *Ccr2*^+/+^ cancer cells, we next examined the rates of proliferation and apoptosis in the tumors collected during the phase when *Ccr2^−/−^* tumors were growth-restricted, 5–6 weeks post-transplantation. There was no difference in proliferation as determined by nuclear Ki67 staining (**Fig. 2c**), and primary *Ccr2*^+/+^ and *Ccr2^−/−^* cancer cells also grew similarly *in vitro* (**Fig. 2d**). However, we found a higher percentage of cancer cells undergoing apoptosis in tumors from *Ccr2^−/−^* cancer cells than from *Ccr2*^+/+^ cancer cells (**Fig. 2e**). Furthermore, *Ccr2^−/−^* cancer cells were more sensitive to serum-free conditions (**Fig. 2f**), consistent with a previous study showing that CCR2 signaling can protect cancer cells from apoptosis induced by serum starvation (Fang et al., 2012).

**Fig. 2:**
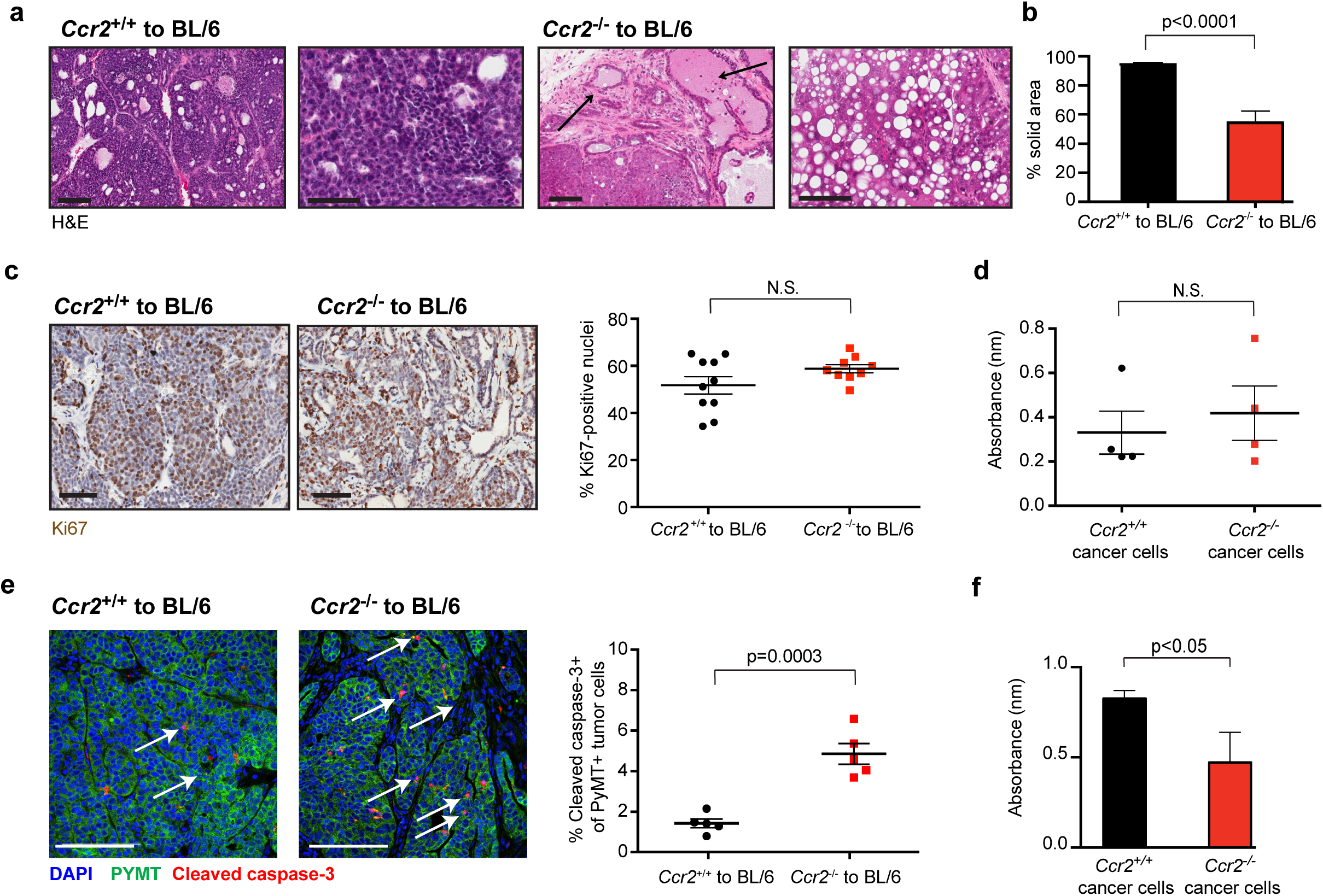
Loss of *Ccr2* increases apoptosis *in vivo* with no effect on proliferation. **a.** Four representative photomicrographs of H&E-stained tumors derived from *Ccr2*^+/+^ and *Ccr2*^−/−^ cancer cells. Arrows denote cystic areas; also note lipid vacuoles in tumors from *Ccr2*^−/−^ cancer cells (Scale bar=100 μm). **b.** Histology score of solid area in tumors from *Ccr2*^+/+^ and *Ccr2*^−/−^ transplants (mean +/− SEM, Student’s t-test; n=10 in *Ccr2*^+/+^ and n=8 in *Ccr2*^−/−^). **c.** Proliferation is unchanged between *Ccr2*^+/+^ and *Ccr2*^−/−^ tumor transplants during the growth-restricted phase, as determined by Ki67-positive nuclear stain. Left panels are representative photomicrographs (Scale bar=100 μm), and right panel shows quantification (mean +/− SEM, Student’s t-test; n=10 and 9 for *Ccr2*^+/+^ and *Ccr2*^−/−^ transplants, respectively). **d.** Proliferation is unchanged in *Ccr2*^+/+^ and *Ccr2*^−/−^ cancer cells *in vitro* during the growth-restricted phase, as determined by absorbance after 24 h using CellTiter 96 AQueous One Solution Cell Proliferation (MTS) Assay (mean +/− SEM, Student’s t-test; n=4). **e.** *Ccr2*^−/−^ cancer cells have an increased apoptotic index compared to *Ccr2*^−/−^ cancer cells during the growth-restricted phase, as determined by double immune staining for PyMT and cleaved caspase-3. Left panels are representative photomicrographs, with arrows indicating apoptotic cancer cells (Scale bar=100 μm); right panel shows quantification (mean +/− SEM, Student’s t-test; n=5). Each dot represents an average of five random fields of views from one tumor. **f.** *Ccr2*^−/−^ cancer cells are less viable than *Ccr2*^+/+^ cancer cells in serum-free conditions, as determined by absorbance in MTS assay after serum starvation for 24 h (mean +/− SEM, Student’s t-test; n=3).

### *Ccr2^−/−^* cancer cells have increased expression of interferon (IFN) response genes and of genes involved in MHC class I antigen presentation

To determine which pathways in the *Ccr2^−/−^* cancer cells led to reduced tumor growth and increased cell death, we isolated cancer cells from *Ccr2*^+/+^ and *Ccr2^−/−^* transplanted tumors formed in wild-type hosts for equal amounts of time and performed transcriptome profiling by RNA-seq. Overall, 520 genes were differentially expressed (adjusted p-value <0.05), ∼40% of which were upregulated in *Ccr2*^−/−^ cancer cells compared to *Ccr2^+/+^* cancer cells. Among the differentially expressed genes, we detected upregulation of MHC class I (H2-K1) and Tap1, two genes required for antigen presentation, and IFI27, an IFN response gene, in the *Ccr2^−/−^* cancer cells (**Fig. 3a**). Vimentin, which is found in poorly differentiated epithelial cells and in mesenchymal cells, was downregulated in *Ccr2^−/−^* cancer cells. This is consistent with a previous study describing a mesenchymal and invasive phenotype in CCR2-expressing cancer cells (Hu et al., 2019). Additionally, gene set enrichment analysis (GSEA) showed robust expression enrichment of the IFN-γ response hallmark gene set, which included genes involved in antigen processing and presentation, in *Ccr2^−/−^* cancer cells compared to *Ccr2*^+/+^ cancer cells (**Fig. 3b**). We also observed a pronounced luminal gene expression signature in the *Ccr2^−/−^* cancer cells (**Fig. 3b**), in agreement with the noted histological differences (**Fig. 2a, b**). Consistently, gene ontology (GO) term analysis revealed altered expression of pathways involved in 1) regulation of the adaptive immune system and in 2) response to immune-mediated cell killing (**Fig. 3c**).

**Fig. 3:**
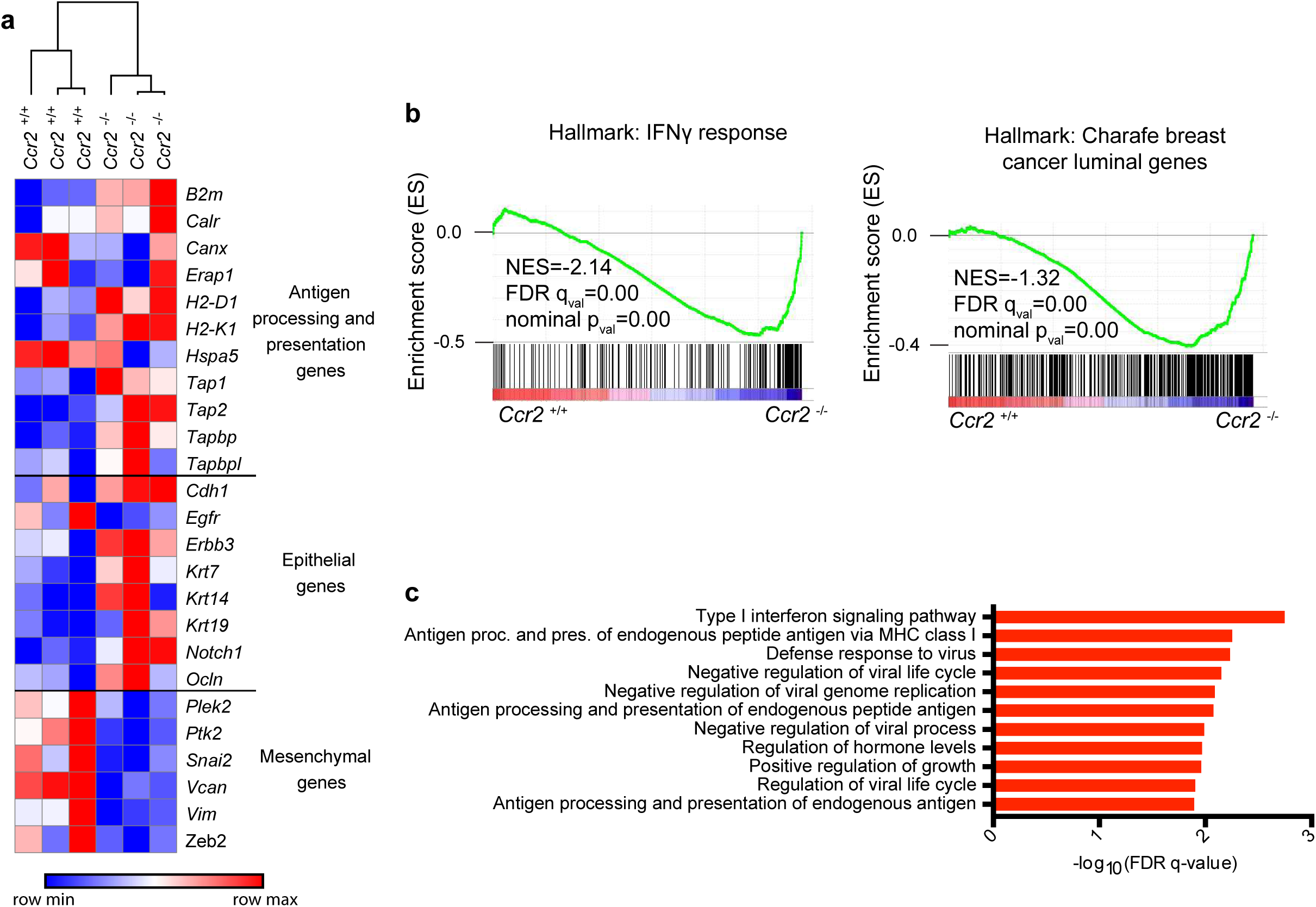
*Ccr2^−/−^* cancer cells have increased expression of interferon (IFN) response genes and of genes involved in MHC class I antigen presentation. **a.** A hierarchically clustered heat map of *Ccr2^+/+^* and *Ccr2^−/−^* cancer cell samples, collected during the growth-restricted phase, based on expression levels of select genes associated with antigen processing and presentation, as well as epithelial-to-mesenchymal transition (n=3 for *Ccr2^+/+^* and *Ccr2^−/−^* tumors). **b.** Gene set enrichment analysis plots showing enrichment of IFN-γ response and luminal gene set expression in *Ccr2^−/−^* cancer cells (NES=normalized enrichment score) **c.** Gene ontology (GO) term enrichment plots showing the most significantly overrepresented GO terms in *Ccr2^−/−^* cancer cells compared to *Ccr2^+/+^* cancer cells.

### Cancer cell CCR2 promotes immune suppression

The increased expression of genes associated with IFN response and antigen presentation, together with the prolonged growth delay and increased apoptosis in tumors from *Ccr2^−/−^* cancer cells, are consistent with an ongoing adaptive immune response in the tumors. We therefore determined whether there were any differences in the infiltrating immune cell populations by flow cytometry (for gating strategy, see **Fig. S2**). There were significantly more CD8+ CTLs and fewer CD4+ helper T cells in tumors from *Ccr2^−/−^* cancer cells than in tumors from *Ccr2*^+/+^ cancer cells (**Fig. 4a**), with no overall change in CD3+ T cell infiltration (**Fig. S3a**). Increased infiltration of CD8+ cells into tumors derived from *Ccr2^−/−^* cancer cells was also evident by immunofluorescence (**Fig. 4b, c**). PD-1 is an inhibitory checkpoint surface molecule expressed by activated CD8+ T cells (Chen et al., 2016a), and its expression was increased on CD8+ T cells from tumors from *Ccr2^−/−^* cancer cells (**Fig. 4d**). CTLs kill, in part, through the release of granzyme B, a process that requires lysosomal-associated membrane protein 1 (LAMP-1, also known as Cluster of Differentiation 107a, CD107a). In tumors from *Ccr2^−/−^* cancer cells, LAMP-1 was increased on the cell surface of CD8+ T cells (**Fig. 4e**), while granzyme B levels were reduced in CD8+ T cells (**Fig. 4f**), indicating increased levels of prior CD8+ degranulation, and therefore cytotoxic activity. Regulatory T lymphocytes (CD4+FoxP3+ regulatory T cells), which are typically immunosuppressive, were more abundant in tumors from *Ccr2^−/−^* cancer cells (**Fig. S3b**).

**Fig. 4:**
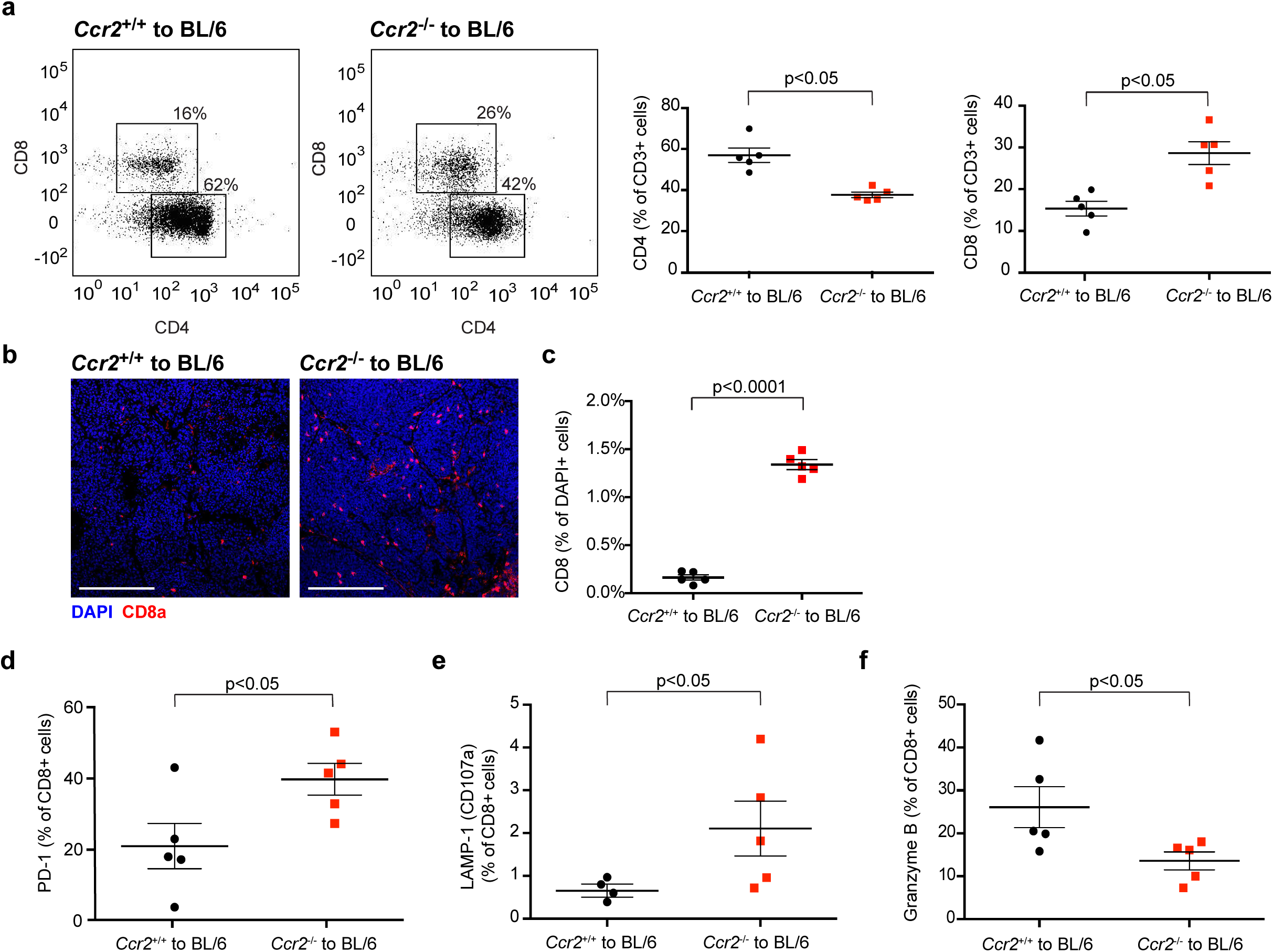
Tumors derived from *Ccr2*^−/−^ cancer cells have increased infiltration of activated CTLs. **a.** Tumors from *Ccr2*^−/−^ cancer cells have fewer CD4+ T cells and more CD8+ T cells during the growth-restricted phase, as determined by flow cytometry gated on CD45+CD3+ cells. Left panels are representative examples of dot plots, and right panels show quantification of CD4+ and CD8+ cells (mean +/− SEM, Student’s t-test; n=5). **b.** Representative immunofluorescence staining showing that more CD8a+ T cells (red) infiltrate into tumors derived from *Ccr2*^−/−^ cancer cells than tumors from *Ccr2^+/+^* cancer cells during the growth-restricted phase (Scale bars=100 μm). **c.** Quantification of CD8a+ T cells in the tumors (mean +/− SEM, Student’s t-test; n=5 tumors. Each dot represents the average of five random fields of views of one tumor). **d.** PD-1 levels were increased in CD8+ T cells in tumors from *Ccr2*^−/−^ cancer cells during the growth-restricted phase, as determined by flow cytometry gating on CD45+CD3+CD8+ cells (mean +/− SEM, Student’s t-test; n=5). **e.** LAMP-1 (CD107a) levels were increased in CD8+ T cells in *Ccr2*^−/−^ tumors during the growth-restricted phase, as determined by flow cytometry gated on CD45+CD3+CD8+ cells (mean +/− SEM, Student’s t-test; n=4 and 5 for *Ccr2*^+/+^ and *Ccr2*^−/−^ transplants, respectively). **f.** Intracellular granzyme B levels were decreased in CD8+ T cells in *Ccr2*^−/−^ tumors during the growth-restricted phase, as determined by flow cytometry gated on CD45+CD3+CD8+ cells, after 2 h of incubation with Brefeldin A (mean +/− SEM, Student’s t-test; n=5).

Tumors from *Ccr2^−/−^* cancer cells had multiple changes in the adaptive immune cell infiltrate, all consistent with an active immune response. Therefore, we tested whether an adaptive immune response contributed to the differences in growth rates between the *Ccr2*^+/+^ and *Ccr2^−/−^* tumors by transplanting cancer cells in parallel into T cell-deficient athymic (nude) or fully immunocompetent mice (**Fig. 5a**). In the immunocompetent hosts, tumors from *Ccr2^−/−^* cancer cells grew significantly slower than tumors from *Ccr2^+/+^* cancer cells (**Fig. 5b**), as previously observed (**Fig. 1g**). In contrast, in the T cell-deficient hosts, the *Ccr2^−/−^* tumors grew at a similar rate as the *Ccr2^+/+^* tumors (**Fig. 5c**). Consistently, tumors derived from *Ccr2^−/−^* cancer cells grew as fast as tumors derived from *Ccr2^+/+^* cancer cells when CD8+ T cells were depleted from wild-type mice (**Fig. S3c**). Furthermore, in the T cell-deficient, athymic hosts, tumors from *Ccr2^−/−^* cancer cells no longer had more apoptotic cells than tumors from *Ccr2*^+/+^ cancer cells (**Fig. 5d**). Interestingly, the differentiated histological phenotype of the tumors from the *Ccr2^−/−^* cancer cells was maintained in the athymic mice (**Fig. 5e, f**). Thus, the histological differences were not the result of an altered adaptive immune response. Together, these data demonstrate that CCR2 expressed by cancer cells enables the tumors to escape the adaptive immune response.

**Fig. 5:**
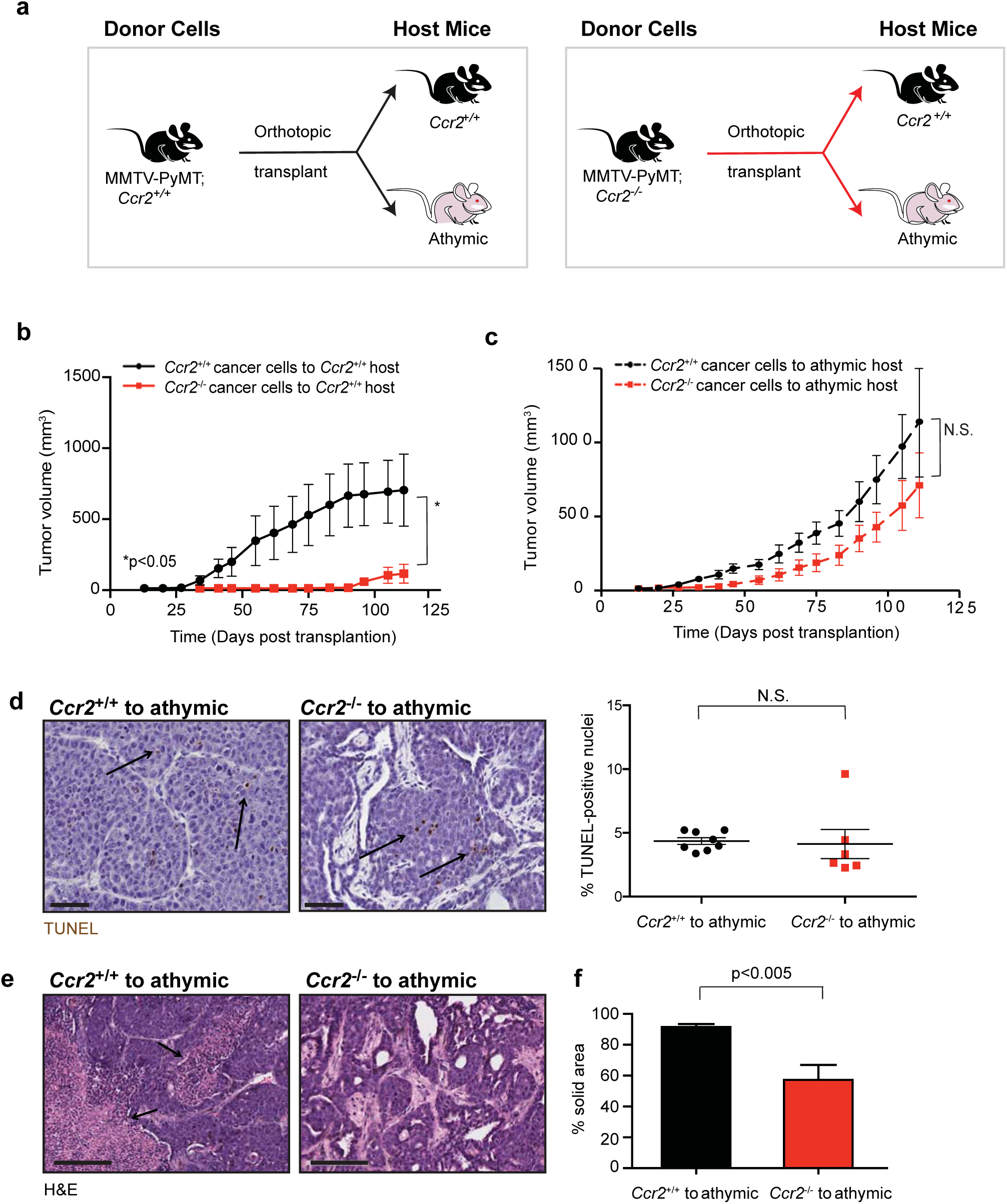
Cancer cell *Ccr2* tumor growth requires functional adaptive immunity. **a.** Schematic of experimental design. **b.** Tumors from *Ccr2*^+/+^ cancer cells grow faster than tumors from *Ccr2*^−/−^ cancer cells in mice with intact immune systems. Tumor burden was determined by weekly caliper measurement. Mice were sacrificed at IACUC-approved endpoint (mean +/− SEM, two-way ANOVA; n=20 per condition). **c.** Tumors from *Ccr2*^+/+^ cancer cells grow similarly to tumors from *Ccr2*^−/−^ cancer cells in athymic host mice. Tumor burden was determined by weekly caliper measurement. Mice were sacrificed at IACUC-approved endpoint (mean +/− SEM, two-way ANOVA; n=20 per condition). **d.** Apoptosis was unchanged between *Ccr2*^+/+^ and *Ccr2*^−/−^ transplants to athymic hosts, as determined by TUNEL stain. Left panels are representative photomicrographs (Scale bar=100 μm), with arrows pointing to apoptotic cells; right panel shows quantification (mean +/− SEM, Student’s t-test; n=8 and 6 for *Ccr2*^+/+^ and *Ccr2*^−/−^ transplants, respectively). **e.** Representative H&E staining of transplanted tumors from *Ccr2*^+/+^ and *Ccr2*^−/−^ cancer cells to nude hosts. Arrows indicate highly necrotic areas (Scale bar=100 μm). **f.** Histology score of solid area of transplanted tumors from *Ccr2*^+/+^ and *Ccr2*^−/−^ cancer cells to athymic hosts (mean +/− SEM, Student’s t-test; n=10 in *Ccr2*^+/+^ and n=9 in *Ccr2*^−/−^).

We next speculated that the reduced growth of tumors from *Ccr2^−/−^* cancer cells could be a result of increased sensitivity to CTL-induced cell death, given that the *Ccr2^−/−^* cancer cells showed increased sensitivity to serum-free conditions and that the tumors grew normally when CD8+ T cells were depleted. To test this possibility, we used the MMTV-PyMT-chOVA model (Engelhardt et al., 2012), which is driven by the PyMT oncogene and co-expresses ovalbumin (OVA) as a model tumor antigen and mCherry for tracking. We generated MMTV-PyMT-chOVA;*Ccr2^+/+^* and MMTV-PyMT-chOVA;*Ccr2^−/−^* mice and found that similar to the MMTV-PyMT;*Ccr2^−/−^* tumors, the MMTV-PyMT-chOVA;*Ccr2^−/−^* tumors grew slower than MMTV-PyMT-chOVA;*Ccr2^+/+^* tumors (**Fig. 6a**). When primary cancer cells were isolated and challenged with activated CTLs isolated from OT-1 mice (where T cells are engineered to recognize an OVA peptide presented by MHC class I), we found that MMTV-PyMT-chOVA;*Ccr2^−/−^* cancer cells were approximately twice as sensitive to CTLs as MMTV-PyMT-chOVA;*Ccr2*^+/+^ cancer cells (**Fig. 6b**). Upon treatment with a pharmacological inhibitor against CCR2 (RS 102895 hydrochloride (Mitchell et al., 2013)), the MMTV-PyMT-chOVA;*Ccr2*^+/+^ cancer cells became as sensitive to the OVA-specific CTLs as MMTV-PyMT-chOVA;*Ccr2^−/−^* cancer cells (**Fig. 6c**). The CCR2 inhibitor also increased specific cytolysis of OVA-expressing E0771 murine breast cancer cell line to the OVA-specific CTLs (**Fig. S3d**). In contrast, the CCR2 inhibitor had no effect on the specific cytotoxicity of the CTLs to the MMTV-PyMT-chOVA;*Ccr2^−/−^* cancer cells (**Fig. 6c**).

**Fig. 6:**
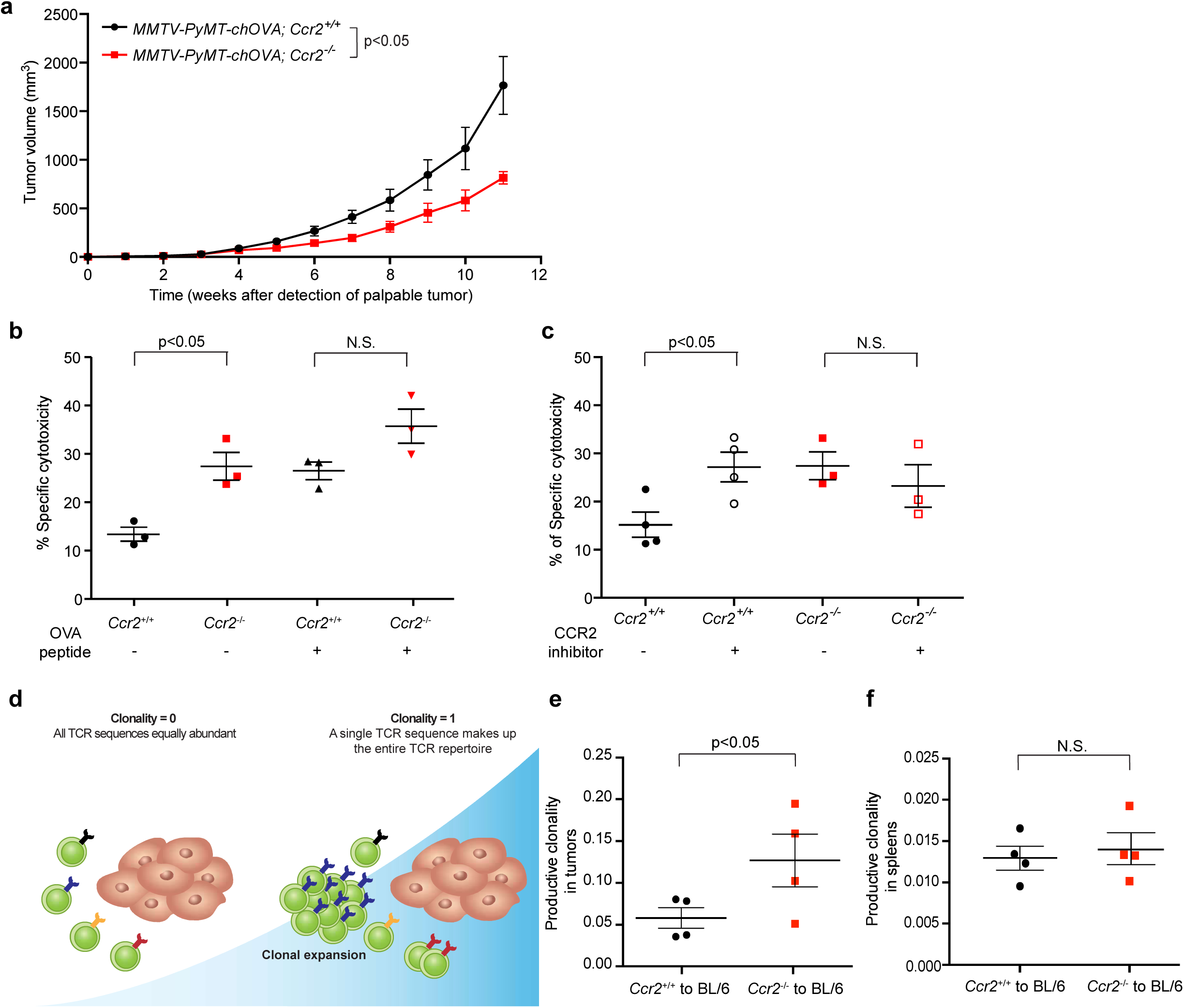
Cancer cell *Ccr2* signaling protects against T cell-mediated killing. **a.** Primary tumor growth is reduced in MMTV-PyMT-chOVA;*Ccr2*^−/−^ mice compared to MMTV-PyMT;*Ccr2*^+/+^ mice, as determined by weekly caliper measurement once the tumors were palpable (mean +/− SEM, two-way ANOVA; n=6 MMTV-PyMT;*Ccr2*^+/+^, and n=5 MMTV-PyMT;*Ccr2*^−/−^ mice). **b.** *Ccr2^−/−^* cancer cells are more sensitive to T cell cytotoxicity than *Ccr2^+/+^* cancer cells. Cytotoxicity of CD8+ OT-1 T cells toward MMTV-PyMT-chOVA;*Ccr2^+/+^* or MMTV-PyMT-chOVA;*Ccr2^−/−^* primary cancer cells, with or without OVA peptides, measured by chromium (Cr^51^) release (mean +/− SEM, Student’s t-test; n=3). **c.** Inhibiting CCR2 increased T cell cytotoxicity against *Ccr2^+/+^* cancer cells. Cytotoxicity of CD8+ OT-1 T cells toward MMTV-PyMT-chOVA;*Ccr2^+/+^* or MMTV-PyMT-chOVA;*Ccr2^−/−^* cancer cells, with or without CCR2 inhibitor, measured by chromium (Cr^51^) release (mean +/− SEM, Student’s t-test; each dot is the average of triplicate from an independent experiment. n=4 and 3 for MMTV-PyMT-chOVA;*Ccr2^+/+^* and MMTV-PyMT-chOVA;*Ccr2^−/−^* cancer cells, respectively). **d.** Model of clonality measured by T cell receptor (TCR) sequencing. **e.** Productive clonality in tumors from *Ccr2*^+/+^ and *Ccr2*^−/−^ cancer cells (mean +/− SEM, Student’s t-test; n=4). **f.** Productive clonality in spleens was unchanged between *Ccr2*^+/+^ and *Ccr2*^−/−^ tumors (mean +/− SEM, Student’s t-test; n=4).

To test if *Ccr2* expression in cancer cells also regulated sensitivity to CTLs *in vivo,* we measured the clonal expansion of T cells, an indicator of active proliferation of antigen-specific T cells, by sequencing the complementarity determining region 3 (CDR3) region of the T cell receptor (TCR) β chain. We found that clonality, a value corresponding to the extent of clonal expansion of T cells (**Fig. 6d**), was significantly higher in tumors from *Ccr2^−/−^* cancer cells than in tumors from *Ccr2*^+/+^ cancer cells (**Fig. 6e**), while there was no difference in clonality in the spleens of these groups of mice (**Fig. 6f**). These results suggested that tumor-recognizing CTLs were actively expanding due to antigen recognition in tumors from *Ccr2^−/−^* cancer cells, but that this expansion did not alter the systemic T cell response. Both *Ccr2*^+/+^ and *Ccr2^−/−^* cancer cells express the PyMT tumor antigen, and *Ccr2*^+/+^ cancer cells formed tumors at the same rate regardless of whether *Ccr2*^+/+^ or *Ccr2^−/−^* cancer cells were transplanted to the contralateral mammary gland (**Fig. S3e**). Together, these results suggest that the expression of CCR2 by cancer cells enables the tumors to establish a locally immune-suppressive microenvironment.

### Cancer cell CCR2 alters MHC class I and PD-L1 expression

CCR2 signaling can activate STAT transcription factors in cancer cells (Chen et al., 2015; Izumi et al., 2013), and altered STAT1 or STAT3 signaling in cancer cells can in turn lead to a less immune-suppressive microenvironment (Ahn et al., 2017; Jones et al., 2016). Therefore, we tested whether STAT activation was different between tumors from *Ccr2^+/+^* and *Ccr2^−/−^* cancer cells. We found no consistent differences between the activation of STAT1, STAT3, or p65/RELA when whole tumor lysate was assayed by Western blot (**Fig. 7a**). By immunohistochemistry, we found that the tumors contained patches of cancer cells with high expression of phopho-STAT1, with more of these patches in the tumors from *Ccr2^−/−^* than from *Ccr2*^+/+^ cancer cells (**Fig. 7b, c**). No difference was observed for phospho-STAT3.

**Fig. 7:**
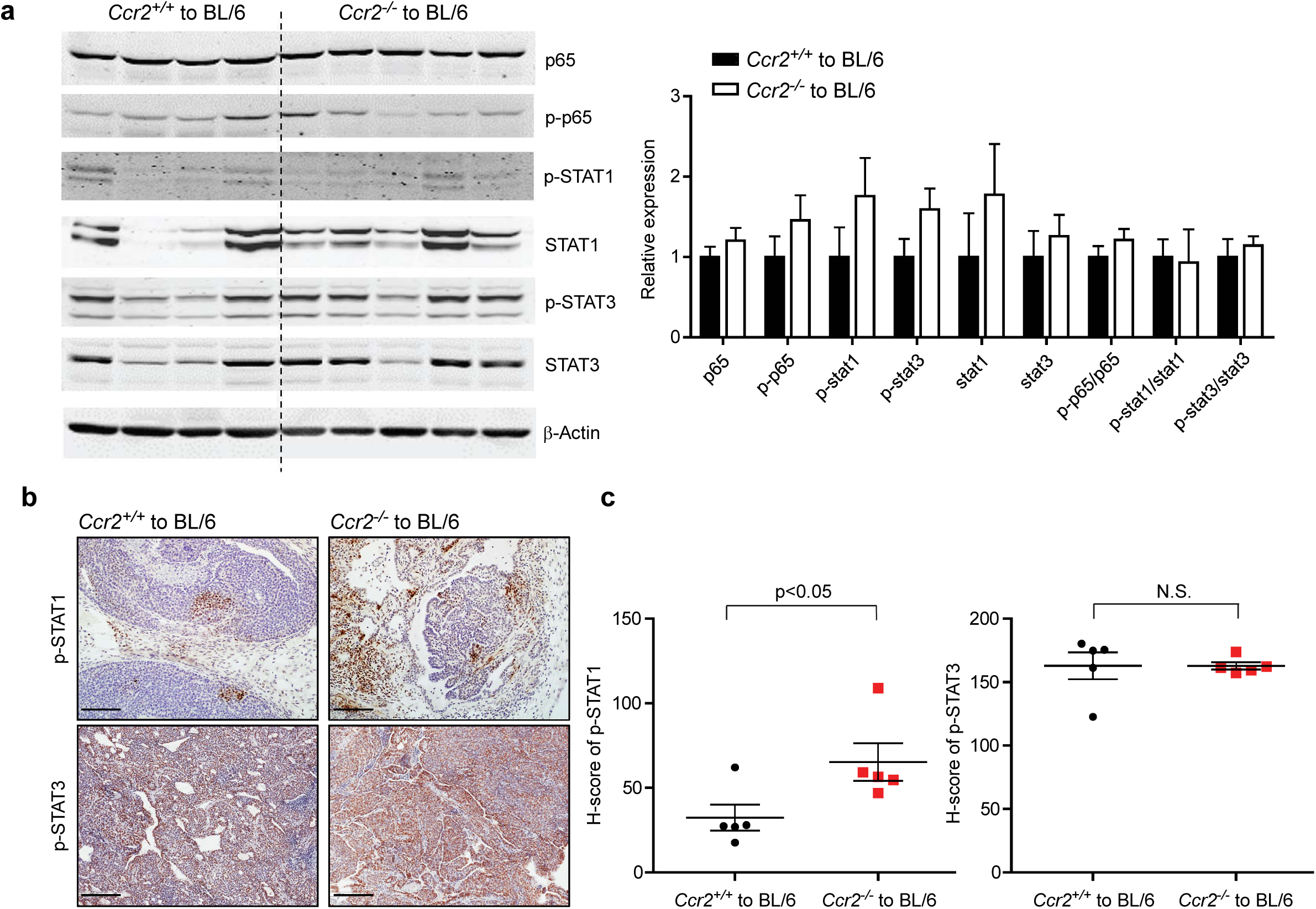
Localized STAT1 activation in *Ccr2^−/−^* tumors. **a.** Comparison of STAT1 activation in tumors from *Ccr2^+/+^* or *Ccr2^−/−^* cancer cells during the growth-restricted phase, by Western blot analysis (left). Each lane contains a protein sample from a tumor from a different mouse. Quantifications of protein expression normalized to β-actin levels (right) (mean +/− SEM, n=4 for *Ccr2^+/+^*, n=5 for *Ccr2^−/−^*). **b.** Representative photomicrographs of p-STAT1 (top) and p-STAT3 (bottom) staining in sections of tumors from *Ccr2^+/+^* or *Ccr2^−/−^* cancer cells. Scale bar=100 μm. **c.** Quantification of the immunohistochemical staining in (b), using H-score, shows higher STAT1 phosphorylation in tumors from *Ccr2^−/−^* than from *Ccr2^+/+^* cancer cells, but similar levels of STAT3 phosphorylation (mean +/− SEM, Student’s t-test; n= 5).

We next sought to understand how CCR2 signaling in cancer cells resulted in more efficient immune suppression. RNA-seq analysis of the cancer cells showed increased expression of genes involved in antigen presentation in the *Ccr2^−/−^* tumors (**Fig. 3a**). Consistently, we found high levels of MHC class I expressed by cancer cells in MMTV-PyMT-chOVA; *Ccr2^−/−^* mice compared to the variable, low to intermediate, levels expressed by cancer cells in MMTV-PyMT-chOVA;*Ccr2^+/+^* mice (**Fig. 8a**). Similarly, when cancer cells were transplanted to immunocompetent host mice, expression of MHC class I was higher by *Ccr2^−/−^* cancer cells than by *Ccr2^+/+^* cancer cells, as shown by immunofluorescence (**Fig. 8b**). However, in T cell-deficient, athymic host mice, MHC class I levels were similar on *Ccr2*^+/+^ and *Ccr2^−/−^* cancer cells (**Fig. 8c**). These data suggest that MHC class I expression is not directly regulated by CCR2 signaling in the cancer cells, but rather is regulated by the different immune responses between these tumors.

**Fig. 8:**
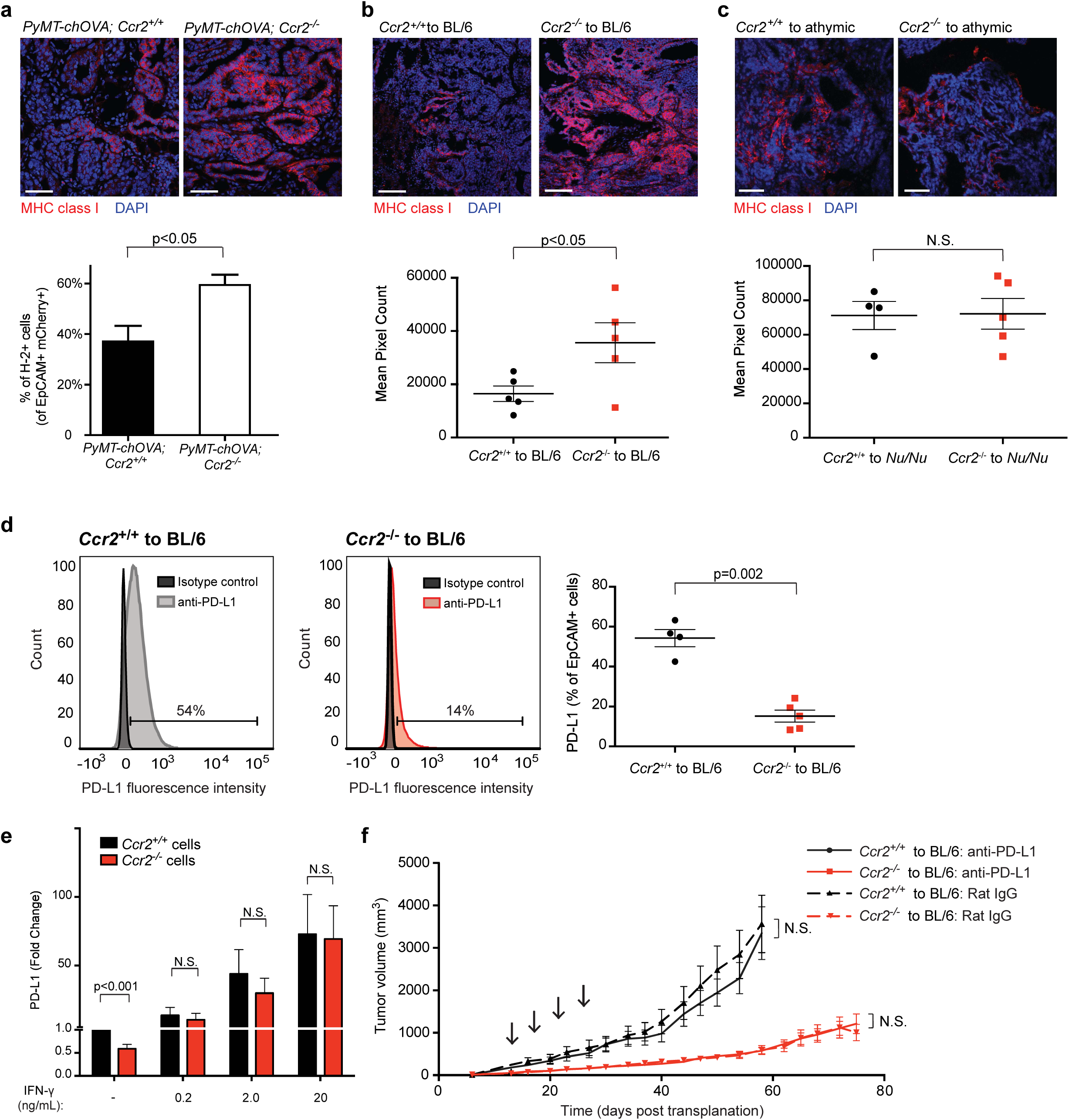
MHC class I expression is reduced and PD-L1 expression increased in *Ccr2^+/+^* cancer cells in immunocompetent mice. **a.** MHC class I expression is increased in MMTV-PyMT-chOVA;*Ccr2*^−/−^ tumors compared to MMTV-PyMT-chOVA;*Ccr2*^+/+^ tumors during the growth-restricted phase, as determined by immunofluorescence staining (upper panels: representative images; Scale bar=50 μm) and by flow cytometry (bottom panel), gated on EpCAM+, mCherry+ cells (mean +/− SEM, Student’s t-test; n=8 and 4 for PyMT-chOVA;*Ccr2*^+/+^ and PyMT-chOVA;*Ccr2*^−/−^ tumors, respectively). **b.** MHC class I is increased on *Ccr2*^−/−^ compared to *Ccr2*^+/+^ cancer cells after transplantation into C57BL/6 hosts during the growth-restricted phase, as determined by immunofluorescence staining (upper panels; Scale bar=50 μm). Quantification of mean pixel count (bottom panel, mean +/− SEM, Student’s t-test; n= 5). **c.** MHC class I is unchanged on *Ccr2*^+/+^ and *Ccr2*^−/−^ cancer cells transplants to athymic hosts, as determined by immunofluorescence staining (upper panels; Scale bar=50 μm). Quantification of mean pixel count (bottom panel, mean +/− SEM, Student’s t-test; n=4 and 5 for *Ccr2*^+/+^ and *Ccr2*^−/−^ transplants, respectively). **d.** PD-L1 is decreased on *Ccr2*^−/−^ cancer cells, as determined by flow cytometry gated on EpCAM+ cells (mean +/− SEM, Student’s t-test; n=5). **e.** PD-L1 mRNA levels are decreased in cultured *Ccr2*^−/−^ cancer cells, but the cells respond equally well to IFN-γ treatment as *Ccr2*^−/−^ cancer cells, as determined by qPCR. *Ccr2*^+/+^ and *Ccr2*^−/−^ cancer cells growing in culture were treated with either 0, 0.2, 2.0, or 20 ng/mL of recombinant IFN-γ for 48 h (mean +/− SEM, Student’s t-test; n=3). **f.** Treatment with four intraperitoneal doses of anti-PD-L1 antibody (200 μg/injection) had no significant effect on growth of *Ccr2*^+/+^ or *Ccr2*^−/−^ tumors *in vivo*, compared to control rat IgG antibody (arrows indicate treatments; mean +/− SEM, two-way ANOVA; n=20 tumors for all conditions).

PD-L1 is a potent suppressor of T cell activation and proliferation and is expressed on many types of cancer cells (Freeman et al., 2000). PD-L1 was significantly higher expressed on *Ccr2*^+/+^ cancer cells than on *Ccr2^−/−^* cancer cells (**Fig. 8d**). Since both MHC class I and PD-L1 are IFN-γ response genes, we compared *Ifn-g* mRNA levels during the early tumor phase (three weeks post-transplantation) and found significantly higher levels of *Ifn-g,* as well as *Cxcl9,* in whole tumor lysate from tumors derived from *Ccr2^-/^*^-^ compared to *Ccr2^+/+^* cancer cells by RT-qPCR (**Fig. S4a**). RNA FISH revealed that the majority of the *Ifn-g* mRNA was expressed by infiltrating CD8+ T cells, and that *Ifn-g* levels were more variable during the growth-restricted phase of tumors than during the early tumor establishment phase (**Fig. S4b, c, d**). *In vitro,* recombinant IFN-γ treatment resulted in similar upregulation of *Pd-l1* mRNA levels in *Ccr2*^+/+^ and *Ccr2^−/−^* cancer cells (**Fig. 8e**). Thus, our data suggest that *Ccr2^−/−^* cancer cells would be able to respond to increased IFN-γ levels in the tumor microenvironment by upregulating PD-L1. Since we see reduced PD-L1 expression on the *Ccr2^−/−^* cancer cells *in vivo* in the context of elevated *Ifn-g,* other factors must repress PD-L1 expression *in vivo*. Lastly, we tested whether PD-L1 blockade would alleviate immune suppression of the MMTV-PyMT tumors *in vivo*. However, anti-PD-L1 therapy had no significant effect on the growth of neither the *Ccr2*^+/+^ tumors nor the low PD-L1-expressing *Ccr2^−/−^* tumors (**Fig. 8f**). Thus, although reduced PD-L1 likely contributed to an increased sensitivity to CTLs in tumors from *Ccr2^−/−^* cancer cells, targeting PD-L1 expression was insufficient to reduce growth of the *Ccr2*^+/+^ tumors.

### Cancer cell CCR2 prevents the infiltration of cross-presenting CD103+ DCs

We next examined the infiltration levels of immune populations that might be involved in immune suppression in the *Ccr2*^+/+^ tumors. There was no difference in CD11b+MHC class II+F4/80+ macrophages between tumors from *Ccr2*^+/+^ and *Ccr2^−/−^* cancer cells (**Fig. S4e**). There were, however, significantly fewer CD11b+MHC class II- myeloid cells in tumors from *Ccr2^−/−^* cancer cells (**Fig. S4f**). This population includes subsets of MDSCs, immature myeloid cells with immune-suppressive activity. Specifically, we found fewer granulocytes and granulocytic MDSCs (CD11b+CD11c-Ly6G+Ly6C^low^ cells) and more inflammatory monocytes and myeloid MDSCs (CD11b+CD11c-Ly6G-Ly6C^high^ cells) in tumors from *Ccr2^−/−^* cancer cells than from *Ccr2*^+/+^ cancer cells (**Fig. S4g, h**).

Since DCs are critical for T cell activation, we performed immunofluorescence for CD11c+, a marker of DCs, and observed more infiltrating CD11c+ cells in the tumors derived from *Ccr2^−/−^* cancer cells than in those derived from *Ccr2*^+/+^ cancer cells (**Fig. 9a**). Furthermore, we found more than double the level of CD103+ DCs—a DC subtype that is very proficient in cross-presenting antigens to CD8+ T cells—in tumors from *Ccr2^−/−^* cancer cells than in those from *Ccr2*^+/+^ cancer cells (**Fig. 9b**). The number of CD103+ DCs was also increased in the draining lymph nodes (dLNs) for these tumors (**Fig. 9c**). A concern when using a transplant system to evaluate immune response is the potential for an immune reaction toward antigens on the transplanted cells. However, there were also more CD103+ DCs in the spontaneously developing tumors in MMTV-PyMT;*Ccr2^−/−^* mice than in MMTV-PyMT;*Ccr2*^+/+^ mice (**Fig. S4i**). Using RT-qPCR from whole tumor lysate and a panel of markers previously shown to be enriched in CD103+ DCs (Broz et al., 2014), we detected more mRNA for *Cd11c* and from activation markers, including *Cd80*, *Cd86*, and *Il12b* (**Fig. 9d**). Consistently, DCs in tumors from *Ccr2^−/−^* cancer cells displayed higher levels of CD86, a surface marker upregulated on activated DCs as determined by flow cytometry (**Fig. 9e**). The difference in CD103+ DC numbers was already apparent in the dLNs two days after injection of irradiated cancer cells into the mammary fat pad (**Fig. 9f**), suggesting a direct signal from the cancer cells. However, infiltration of CD103+ DCs was not increased in tumors from *Ccr2^−/−^* cancer cells in T cell-deficient, athymic mice (**Fig. 9g**). This finding suggests that a feedback loop between T cells and DCs is critical for CD103+ DC recruitment to the tumors.

**Fig. 9:**
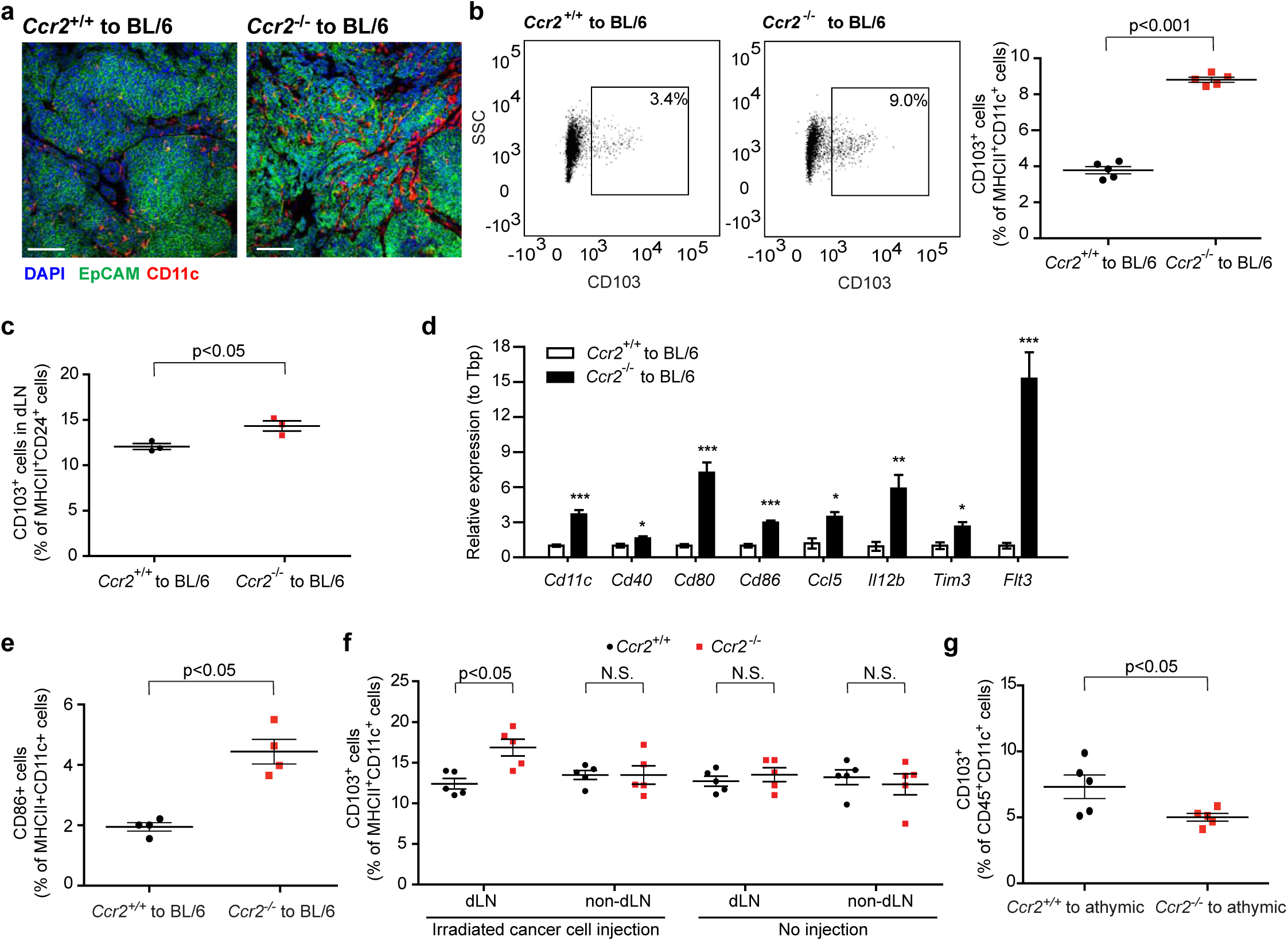
*Ccr2^−/−^* tumors promote infiltration and activation of cross-presenting CD103+ DCs. **a.** Representative immunofluorescence images showing that more CD11c+ dendritic cells (red) infiltrate into *Ccr2*^−/−^ tumors compared to *Ccr2*^+/+^ tumors during the growth-restricted phase (EpCAM+, green; Scale bar=100 μm). **b.** CD103+ DCs are increased in *Ccr2*^−/−^ tumors compared to *Ccr2*^+/+^ tumors during the growth-restricted phase, as determined by flow cytometry gated on CD45+CD11c+MHC class II+ cells. SSC=side scatter (mean +/− SEM, Student’s t-test; n=5). **c.** CD103+ DCs are increased in the draining lymph nodes (dLNs) of *Ccr2*^−/−^ tumors compared to the dLNs of *Ccr2*^+/+^ tumors during the growth-restricted phase, as determined by flow cytometry gated on CD45+CD11c+MHC class II+ cells (mean +/− SEM, Student’s t-test; n=5). **d.** Tumors from transplanted *Ccr2*^−/−^ cancer cells have increased DC infiltration and elevated markers of DC activation compared to tumors from *Ccr2*^+/+^ cancer cells. RT-qPCR on tumors during the growth-restricted phase (mean +/− SEM, Student’s t-test; for *Cd11c*, n=6 in *Ccr2*^+/+^ and n=8 in *Ccr2*^−/−^, for other genes, n=5 in *Ccr2*^+/+^ and n=6 in *Ccr2*^−/−^). **e.** DCs in *Ccr2*^−/−^ tumors are more activated during the growth-restricted phase, as determined by flow cytometry for CD86+ gated on CD45+CD11c+MHC class II+ cells (mean +/− SEM, Student’s t-test; n=5). **f.** Flow cytometry on dLNs with or without injection of irradiated cancer cells into the mammary gland. Note that CD103+ DCs are only increased in the dLNs of the glands after injection of *Ccr2^−/−^* cancer cells, not in non-dLNs from axillary glands (mean +/− SEM, Student’s t-test; n=5). **g.** CD103+ DCs are decreased in *Ccr2*^−/−^ tumors compared to *Ccr2*^+/+^ tumors in athymic hosts, as determined by flow cytometry gated on CD45+CD11c+ cells (mean +/− SEM, Student’s t-test; n=5).

We next asked whether a factor secreted by *Ccr2^−/−^* cancer cells was responsible for DC maturation toward the CD103+ DC subtype. Indeed, when bone marrow-derived cells were cultured with conditioned medium from *Ccr2^−/−^* or *Ccr2*^+/+^ cancer cells, the total number of DCs was not different, but a significantly higher percentage of CD11c+ cells were positive for CD103+ using conditioned medium from the *Ccr2^−/−^* cells (**Fig. 10a**). Furthermore, the percentage of CD103+CD11c+ cells also increased when using conditioned medium from *Ccr2*^+/+^ cancer cells that had been treated with the CCR2 inhibitor (**Fig. 10b**), whereas adding the CCR2 inhibitor directly to the bone marrow-derived cultures did not by itself increase the percentage of CD103+ cells. Flt3 ligand and granulocyte-macrophage colony-stimulating factor (GM-CSF) can both stimulate DC maturation toward the CD103+ subtype (Broz et al., 2014). Consistently, we found more mRNA for both *Flt3l* and *Csf2* (coding for GM-CSF) in the tumors derived from *Ccr2^−/−^* cancer cells than from *Ccr2*^+/+^ cancer cells by RT-qPCR (**Fig. 10c**). Protein arrays showed that the levels of two CCR2 ligands, CCL2 and CCL12, and most other cytokines were similar between MMTV-PyMT;*Ccr2^+/+^* and MMTV-PyMT;*Ccr2^−/−^* tumors (**Fig. S5a**), while IL-16 and granulocyte colony-stimulating factor (G-CSF) were significantly lower in the MMTV-PyMT;*Ccr2^−/−^* tumors at the protein level. When secretions from overnight cultures of the cancer cells under serum-free conditions were analyzed, CCL2, CCL12, G-CSF, and IL-16 were secreted to the same degree by *Ccr2*^+/+^ and *Ccr2^−/−^* cancer cells (**Fig. S5b**). Lastly, to test the importance of the CD103+ DCs in regulating an immune-controlling microenvironment, we transplanted *Ccr2^−/−^* and *Ccr2*^+/+^ cancer cells into *Batf3^−/−^* mice, which lack the basic leucine zipper transcription factor ATF-like 3 (BATF3) critical for CD103+ DC maturation. In these mice, the growth of *Ccr2^−/−^* cancer cell-derived tumors was restored to that of the *Ccr2*^+/+^ cancer cell-derived tumors (**Fig. 10d**), and as expected, the percentage of infiltrating CD103+ DCs was significantly reduced (**Fig. 10e**). Furthermore, CD8+ T cell infiltration, as well as expression of CD107a and PD-1 by the T cells, were reduced in tumors derived from *Ccr2^−/−^* cancer cells transplanted into *Batf3^−/−^* mice (**Fig. 10f, g, h**). These findings support the notion that CD103+ DC and CD8+ T cells act together to regulate tumor immunity.

**Fig. 10:**
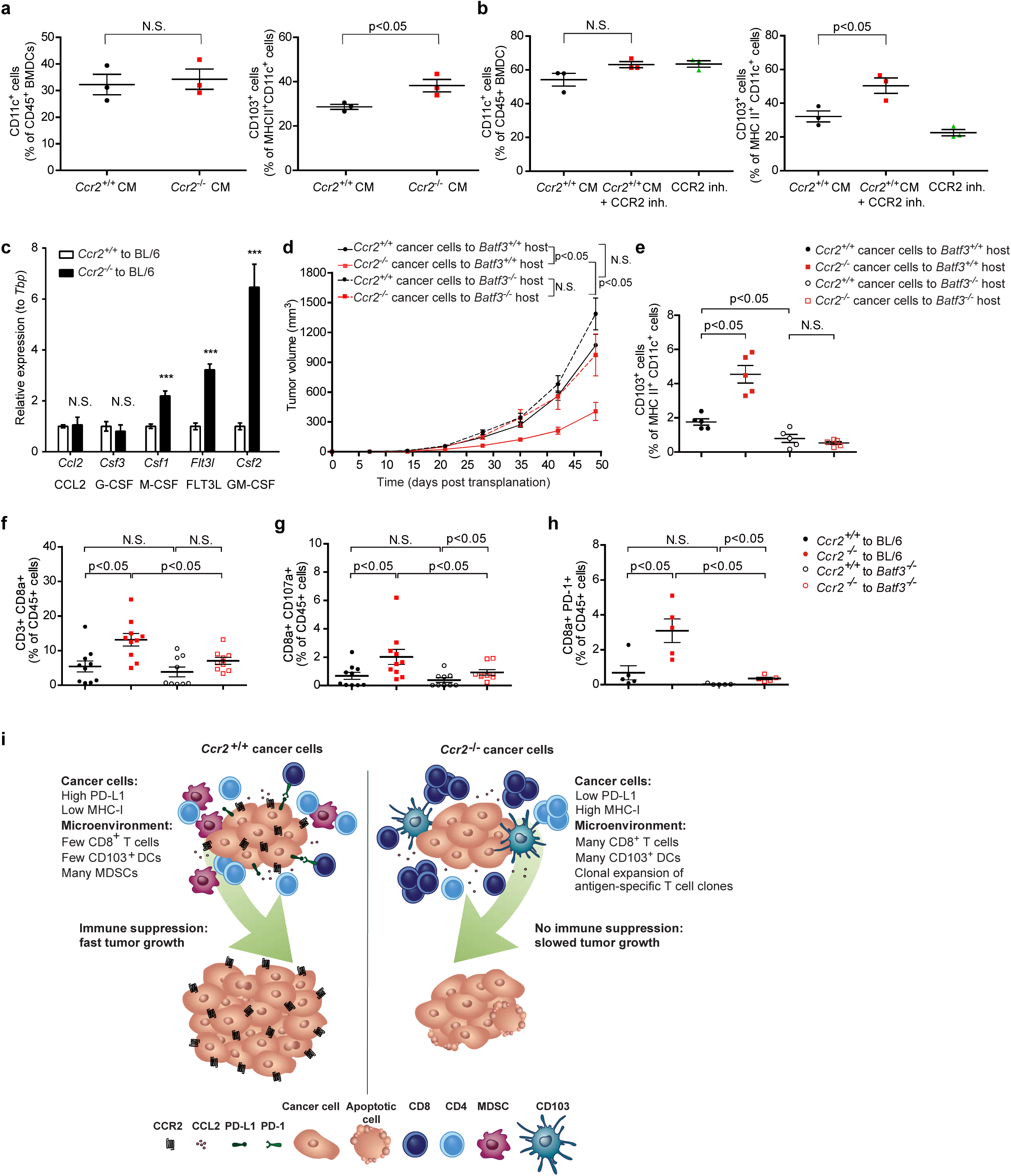
Cross-presenting CD103+ DCs are critical for immune control of *Ccr2^−/−^* cancer cells. **a.** Conditioned medium from *Ccr2*^−/−^ cancer cells induce similar numbers of CD11c DCs (left), but more CD103+ BMDCs (right) *in vitro*, compared to conditioned medium from *Ccr2*^+/+^ cancer cells (mean +/− SEM, Student’s t-test; n=3). **b.** Conditioned medium from *Ccr2*^+/+^ cancer cells cultured with CCR2 inhibitor induced similar numbers of CD11c DCs (left), but more CD103+ BMDCs (right) *in vitro* than conditioned medium from control cells cultured in the absence of the inhibitor medium (mean +/− SEM, Student’s t-test; n=3). **c.** Tumors from transplanted *Ccr2*^−/−^ cancer cells have elevated *Csf2* mRNA (coding GM-CSF) and *Flt3l* (coding FLT3 ligand), but not *Ccl2* or *Csf3* compared to tumors from *Ccr2*^+/+^ cancer cells. RT-qPCR on tumors from transplanted *Ccr2*^+/+^ and *Ccr2*^−/−^ cancer cells during the growth-restricted phase (mean +/− SEM, Student’s t-test; n=5–6 for *Ccr2*^+/+^ tumors and n=6 for *Ccr2*^−/−^ tumors). **d.** Tumor growth is increased in *Batf3*^−/−^ hosts transplanted with *Ccr2*^−/−^ cancer cells, as determined by weekly caliper measurement (mean +/− SEM, two-way ANOVA; n=9 in *Ccr2*^+/+^ to *Batf3*^−/−^ hosts group, and n=10 for the other three groups). **e.** CD103+ DCs are increased in *Ccr2*^−/−^ tumors compared to *Ccr2*^+/+^ tumors in C57BL/6 hosts but are low in *Batf3^−/−^* hosts (mean +/− SEM, Student’s t-tests; n=5). **f.** CD8a+ T cells are increased in early-stage tumors (three weeks) from transplanted *Ccr2*^−/−^ cancer cells compared to tumors from *Ccr2*^+/+^ cancer cells in C57BL/6 hosts, but are low in *Batf3^−/−^* hosts (mean +/− SEM, Student’s t-tests; n=9–10). **g.** Lamp-1 (CD107a) and CD8a double-positive T cells are increased in early-stage tumors (three weeks) from transplanted *Ccr2*^−/−^ cancer cells compared to tumors from *Ccr2*^+/+^ cancer cells, but are low in *Batf3^−/−^* hosts (mean +/− SEM, Student’s t-tests; n=9–10). **h.** PD-1 and CD8a double-positive T cells are increased in early-stage tumors (three weeks) from transplanted *Ccr2*^−/−^ cancer cells compared to tumors from *Ccr2*^+/+^ cancer cells in C57BL/6 hosts, but are low in *Batf3^−/−^* hosts (mean +/− SEM, Student’s t-tests; n=5). **i.** Model of how cancer cell CCR2 orchestrates immune suppression.

## DISCUSSION

The tumor-promoting effects of CCR2 are well-documented. However, they have mostly been attributed to CCR2-expressing immune and other host cells (Lim et al., 2016; Nakasone et al., 2012; Qian et al., 2009; Qian et al., 2011; Wolf et al., 2012). Yet, CCR2 is also upregulated on breast cancer cells in a subset of breast cancers and this upregulation has been correlated with shortened survival (Labovsky et al., 2017). The expression of CCR2 by cancer cells has been proposed to promote tumor growth through increased motility and invasion, cell proliferation, and cell survival (Fang et al., 2012; Hu et al., 2019; Lu et al., 2006). Here, we show that in the MMTV-PyMT mouse model, CCR2 signaling in cancer cells helps breast tumors establish a locally immune-suppressive microenvironment. Compared to tumors from *Ccr2^+/+^* cancer cells, the tumors from *Ccr2^−/−^* cancer cells have higher infiltration of CD103+ DCs and activated CD8+ T cells, as well as lower infiltration of granulocytic MDSCs. Furthermore, the *Ccr2^−/−^* cancer cells have higher surface levels of MHC class I and lower levels of PD-L1. It is likely the combination of all of these changes that results in more effective immune surveillance of the *Ccr2^−/−^* cancer cells and reduced growth of tumors from these cells (**Fig. 10i**). To our knowledge, the ability of cancer cell CCR2 signaling to orchestrate tumor immune suppression has not previously been recognized.

Among the most notable changes in the tumor immune microenvironment caused by targeting cancer cell *Ccr2* were the increased levels of both CD103+ DCs and CD8+ T cells in tumors, and the infiltration of these two cell populations was apparently connected. In athymic mice lacking mature CD8+ cells, the CD103+ DC infiltration was no longer elevated in tumors derived from *Ccr2^−/−^* cancer cells compared to those from *Ccr2^+/+^* cancer cells. Conversely, in *Batf3^−/−^* mice, which lack the transcription factor BATF3 required for CD103+ DC development, CD8+ T cell infiltration was equally low for tumors from *Ccr2^+/+^* and *Ccr2^−/−^* cancer cells. These observations are consistent with several other recent reports intimately linking the function of CD103+ DCs and CD8+ T cells in tumors. First, CD103+ DCs have been shown to secrete chemokines mediating T cell infiltration, including CXCL9 and -10 (de Mingo Pulido et al., 2018; Spranger et al., 2017). Second, secretion of IL-12 from CD103+ DCs has been shown to stimulate CD8+ T cell function (Ruffell et al., 2014). Our data support these previous studies, showing not only a link between CD103+ DC and CD8+ T cell infiltration but also increased *Cxcl9* and *Il-12b* mRNA in tumors derived from *Ccr2^−/−^* cancer cells. Altogether, these data support previous work showing that the regulation of CD103+ DC infiltration is a critical step in tumor immune surveillance (Broz et al., 2014; Meyer et al., 2018; Roberts et al., 2016).

Our data suggest that CCR2 signaling regulates the secretion of a factor(s) from the cancer cells that affects the polarization of DCs to the CD103+ subtype. Several factors have been proposed to induce the polarization of CD103+ DCs, including Flt3L and GM-CSF (Mayer et al., 2014), and consistently, mRNA levels for both of these factors were higher in tumors from *Ccr2^−/−^* cancer cells than from *Ccr2^+/+^* cancer cells. However, it is unclear if Flt3L and GM-CSF could be solely responsible for the difference in CD103+ DC infiltration between tumors from *Ccr2^+/+^* or from *Ccr2^−/−^* cancer cells. A recent study reported that G-CSF secreted from MMTV-PyMT cancer cells can inhibit the differentiation of CD103+ DCs by downregulating interferon regulatory factor-8 in DC progenitors (Meyer et al., 2018). Interestingly, G-CSF was reduced in MMTV-PyMT;*Ccr2^−/−^* tumors compared to MMTV-PyMT;*Ccr2^+/+^* tumors, although secretion of G-CSF from isolated cancer cells was not reduced, suggesting that altered secretion is not a direct effect of CCR2 signaling in cancer cells.

*Ccr2^−/−^* cancer cells had distinct expression of a several genes, including many that were consistent with an IFN response. Among these, increased surface expression of MHC class I might explain why *Ccr2^−/−^* cancer cells are more sensitive to T cell-mediated killing than *Ccr2*^+/+^ cancer cells. However, since *Ccr2*^+/+^ cancer cells did not downregulate MHC class I in T cell-deficient, athymic mice, we speculate that MHC class I expression is not directly regulated by CCR2 signaling in the cancer cells. Rather, the altered expression of IFN response genes, including MHC class I, may be an indirect result of an immune response in the tumors. Altogether, we propose that the cancer cells act on CD103+ DCs, leading to a feedback loop between CD103+ DCs and CD8+ T cells—two immune cell types with highly interconnected functions.

Our study has several translational implications. First, our findings provide a possible explanation as to why CCR2 overexpression in breast cancer cells is associated with shortened survival time (Labovsky et al., 2017). Second, results from an open-label, non-randomized phase I trial using a CCR2 inhibitor against locally invasive pancreatic cancer suggest that the inhibitor enhanced chemotherapy response, reduced macrophage infiltration, and induced an anti-tumor adaptive immune response (Nywening et al., 2016). Our data suggest that the potential clinical benefits of CCR2 inhibitors might be caused by the combined inhibition of CCR2 signaling in both stromal and cancer cells. Interestingly, the loss of even one *Ccr2* allele was sufficient to reduce tumor growth in our study, suggesting that partial inhibition of CCR2 could result in clinical benefit. It is unclear why *Ccr2^+/−^* cancer cells phenocopy the *Ccr2^−/−^* cancer cells, but this phenomenon could be due to reduced dimerization of CCR2 in *Ccr2^+/−^* cancer cells, consistent with a prior report (Rodriguez-Frade et al., 1999), or because surpassing a threshold level of CCR2 signaling is required to induce immune suppression. Finally, recombinant G-CSF and GM-CSF are routinely given with certain types of chemotherapy to support the bone marrow and reduce the risk of lethal febrile neutropenia (Mehta et al., 2015). Our data support prior studies suggesting that these cytokines play complex roles in regulating the immune response against tumors: they appear to have opposing effects on CD103+ DC differentiation, with GM-CSF stimulating CD103+ DC differentiation (Greter et al., 2012; Mayer et al., 2014) and G-CSF inhibiting it (Meyer et al., 2018). Although GM-CSF secretion in the tumor microenvironment potentially enhances immune responses against tumors by stimulating CD103+ DC differentiation, this cytokine can also mobilize immune-suppressive MDSCs, leading to suppression of an anti-tumor immune response (Bayne et al., 2012; Pylayeva-Gupta et al., 2012). Therefore, exploring other approaches than administration of recombinant GM-CSF and G-CSF to specifically boost CD103+ DC maturation could be an important future translational direction, and determining the exact mechanism that causes inhibition of cancer cell CCR2 signaling to promote CD103+ maturation may provide insights into how this process occurs.

We found that tumor onset was similar between MMTV-PyMT;*Ccr2^+/+^* and MMTV-PyMT;*Ccr2^−/−^* mice, while tumor growth was slower. These data are very similar to a prior report studying tumor growth in MTMV-Neu mice, where mice null for *Ccl2*, the ligand for CCR2, or mice treated with a pharmacological CCR2 inhibitor also had similar tumor onset yet reduced tumor growth rates, as did mice treated with a CCR2 inhibitor (Chen et al., 2016b). However, unexpectedly, MMTV-Neu mice lacking *Ccr2* had accelerated, not reduced, tumor growth in this study, and the authors speculated that this could be due to abnormal differentiation of the monocytes in these mice or unintentional effects of the genetic targeting of *Ccr2*. There are several possible explanations for the difference between our result using MMTV-PyMT;*Ccr2^−/−^* mice and that of the MMTV-Neu;*Ccr2^−/−^* mice in the previous publication. First, tumors of MMTV-PyMT mice have more stromal components than tumors of MMTV-Neu mice. Second, the genetic background of the mice used in our experiments was C57BL/6, while it was BALB/c in the case of the MMTV-Neu model. Finally, our tumor transplantation experiments strongly suggest that the tumor growth phenotype in our model is caused by CCR2 expression by cancer cells, but it is not clear whether cancer cells in the MMTV-Neu model express CCR2 or whether the phenotype is due to effects from stromal cells.

Despite the data suggesting that CCR2 inhibition has beneficial effects in cancer treatment, some studies have raised serious concerns about targeting the CCR2 ligand. Cessation of anti-CCL2 therapy resulted in increased pulmonary metastasis and decreased survival in a murine model of breast cancer, due to rapid mobilization of CCR2-expressing monocytes into circulation immediately upon cessation of therapy (Bonapace et al., 2014). Therefore, CCR2 inhibition likely will have to be used in combination with other therapies to maximize the eradication of existing cancer cells and to avoid complications of discontinuing the inhibition. In summary, the divergent results across mouse models, genetic backgrounds, and variable experimental approaches to targeting CCL2 or CCR2 suggest that the effects of targeting this axis can depend on tumor subtype, genetic background, and cell types.

Our findings support the notion that the CCL2/CCR2 axis is an immune modulatory pathway in cancer, but with the added layer of complexity that cancer cells can hijack this chemokine receptor pathway to orchestrate immune suppression. In this study, we have focused on characterizing the cellular mechanisms responsible for the novel link between cancer cell CCR2 signaling and tumor immune suppression. Preclinical experiments suggest that CCR2 targeting strategies can have detrimental effects, including the promotion of metastasis (Bonapace et al., 2014; Li et al., 2013). Therefore, future work that identifies the precise mechanisms by which CCR2 signaling in cancer cells induces immune suppression in breast cancer may set the stage for developing novel immune modulatory therapies. There are ongoing clinical trials using pharmacological inhibitors of CCR2 in cancer patients, and our data suggest that the potential effects of inhibiting CCR2 signaling in cancer cells should be taken into account when interpreting the results from these trials.

## MATERIALS AND METHODS

### Mice

*Ccr2^-^*^/-^, *Batf3^−/−^*, and OT-1 mice (all on a C57BL/6 background) were obtained from Jackson Laboratory (Bar Harbor, ME) and *athymic (Nu/Nu)* mice were obtained from Charles River Laboratory (Wilmington, MA). MMTV-PyMT mice (on C57BL/6 [referred to as BL/6] background) were bred at Cold Spring Harbor Laboratory (CSHL), and MMTV-PyMT-chOVA mice (Engelhardt et al., 2012) were a kind gift from Dr. Mathew Krummel (University of California, San Francisco). MMTV-PyMT and PyMT-chOVA mice were intercrossed with *Ccr2^-^*^/-^ mice. MMTV-PyMT;*Ccr2*^+/+^ and MMTV-PyMT;*Ccr2*^−/−^ mice, and similarly MMTV-PyMT-chOVA;*Ccr2*^+/+^ and MMTV-PyMT-chOVA;*Ccr2*^−/−^ mice, were from the same mouse colony (mice were littermates or their parents were littermates). All the BL/6 host mice were purchased from Jackson Laboratory. All animal experiments were approved by the Institutional Animal Care and Use Committee (IACUC) at CSHL and were conducted in accordance with the NIH Guide for the Care and Use of Laboratory Animals.

### Measurement of *in vivo* tumor growth and lung metastasis

Tumor onset was determined by weekly palpation. Once a tumor was detected, its size was measured weekly by caliper, and tumor volume was calculated as (length x width^2^)/2. For primary tumor growth, mice were either sacrificed at end point, when tumors reached 20 mm or ulcerated (whichever came first) or at indicated time points (early phase: three weeks after transplantation; growth-restricted phase: 5–6 weeks after transplantation). For the lung metastasis assay, mice were sacrificed at either of two IACUC-approved endpoints (when tumors reached 25 mm or ulcerated). To determine metastatic burden, we adapted a previously published method (Nielsen et al., 2001). Briefly, lungs were placed in 4% paraformaldehyde (PFA) in phosphate-buffered saline (PBS) in a vacuum desiccator for 1 h. The lungs remained in 4% PFA at 4°C for 48 h, followed by incubation in 20% sucrose in PBS at 4°C for 48 h. Lungs were embedded in Tissue-Tek O.C.T. compound embedding solution (Sakura, Torrance, CA). The blocks were then placed in a custom-made cutting chamber with razor blade inserts every 2 mm, and the block was cut into 2 mm sections. These sectioned lung pieces were then re-embedded in fresh O.C.T. to allow for cross-sectional cuts from the entire lung tissue. These cross-sections of the lungs were stained with H&E and scanned by an Aperio ScanScope® CS System (Leica Biosystems, Buffalo Grove, IL). The metastatic burden was calculated as the percentage of metastasis/lung area and the number of foci/lung area, determined using Aperio eSlide Capture Devices software (Leica Biosystems). Histology was evaluated by pathologist J.E.W. and scored as percentage solid areas of tumors.

### Primary cancer cell isolation, culture, and transplantation

Aged-matched virgin females (C57BL/6, *Ccr2*^−/−^, *Batf3^−/−^,* or *Nu/Nu*), 6-12 weeks of age, were used as hosts for transplantation. Cancer cells were isolated from 2-3 tumors, each one 8-10 mm in diameter, from MMTV-PyMT;*Ccr2*^+/+^ or MMTV-PyMT;*Ccr2*^−/−^ mice. Tumors were mechanically dissociated and digested for 1 h with 1x collagenase/hyaluronidase (10X Solution, Stem Cell Technology, Cambridge, MA) containing DNase I (2 U/ml, Roche, Pleasanton, CA) in Roswell Park Memorial Institute (RPMI)-1640 medium supplemented with 5% FBS (VWR Life Science Seradigm, Philadelphia, PA). Single cells and debris were removed from the resulting carcinoma organoid preparation by pulse centrifugation in HBSS supplemented with 5% FBS. Purified carcinoma organoids were dissociated into single cell suspension in 0.05% trypsin with 0.1% EDTA supplemented with 2 U/ml of DNase I (Roche) for 2-3 min. Single tumor cells were passed through a 100 μm cell strainer (BD Biosciences, San Jose, CA) and either plated in RPMI-1640 supplemented with 10% FBS for *in vitro* experiments or washed with PBS and immediately injected into the inguinal mammary glands of host mice (2.5×10^5^ in 20 μl of 1:1 PBS/Matrigel; Corning, Corning, NY).

To evaluate the effect of cancer cell *Ccr2* on the anti-tumor immune response locally vs. systemically, contralateral transplantation was conducted with cancer cells with different *Ccr2* genotypes transplanted into each inguinal mammary gland (2.5×10^5^ in 20 μl of 1:1 PBS:Matrigel; Corning, Corning, NY). Cancer cells were isolated as described above and both C57BL/6 and *Ccr2*^−/−^ mice were used as hosts.

To transplant irradiated cancer cells, freshly isolated primary MMTV-PyMT;*Ccr2*^+/+^ or MMTV-PyMT;*Ccr2*^−/−^ cancer cells were plated in a 10 cm petri dish (2×10^6^ per dish) in RPMI with 10% FBS, and received 80 Gy of irradiation in the Gammacell 40 Exactor (Best Theratronics, Ottawa, Ontario, Canada). After irradiation, cancer cells were washed with PBS and injected into the inguinal mammary glands of host mice (2.5×10^5^ in 20 μl of PBS). To generate cancer cell conditioned medium, primary cancer cells from either MMTV-PyMT;*Ccr2*^+/+^ or MMTV-PyMT;*Ccr2*^−/−^ tumors were plated in a 10 cm dish (2×10^6^ per dish) in DMEM/F12 medium plus 10% FBS, 1% penicillin/streptomycin, with or without CCR2 antagonist (RS504393, 10 μM, Sigma-Aldrich, St. Louis, MO) for two days. The conditioned medium was collected and centrifuged at 300 x g for 10 min, and the supernatant was used in experiments.

### *In vivo* antibody treatment

Tumor cells from MMTV-PyMT;*Ccr2*^+/+^ or MMTV-PyMT;*Ccr2*^−/−^ mice were transplanted as described above. For anti-PD-L1 treatment, on days 15, 18, 21, and 24 after transplantation (after tumors had formed), mice received 200 μg of anti-PD-L1 antibody by intraperitoneal injection (Clone 10F.9G2; Bio X Cell, West Lebanon, NH) or control rat IgG2b antibody (Bio X Cell) (Winograd et al., 2015). For CD8+ T cell depletion, mice were injected intraperitoneally with 200 μg of anti-CD8a antibody (Clone 2.43; BE0061; Bio X Cell, West Lebanon, NH) or control rat IgG2b antibody (Clone LTF-2; BE0090; Bio X Cell, West Lebanon, NH) every three days starting from day 0 and until day 30. Tumor size was measured biweekly, and mice were sacrificed at IACUC-approved endpoint.

### Flow cytometry

Tumors were harvested and mechanically dissociated for 30 min with collagenase D (2 mg/ml; Sigma-Aldrich) and DNase I (4 U/ml; Roche) in RPMI. For flow cytometry of cells from lymph nodes, the lymph nodes were forced through a 100 μm cell strainer (BD Biosciences), and flow-through cells were collected for experiments. Cells were resuspended in 1 x HBSS supplemented with 0.5% BSA and centrifuged at 300 x g. The cell suspension was filtered through a 70 μm cell strainer, and red blood cells were lysed using red blood cell lysing buffer (Sigma-Aldrich) for 1 min at room temperature.

For flow cytometry staining, cells (1×10^6^) were incubated with mouse Fc Block (clone 2.4G2, BD Biosciences) for 10 min on ice, and then stained with the appropriate antibodies to surface markers at 4°C for 30 min in the dark or permeabilized and stained with intracellular antibodies overnight (see below for antibodies). Cell viability stain Zombie Red (BioLegend, San Diego, CA) was used to differentiate between dead and live cells. The stained populations were analyzed using an LSR II flow cytometer (BD, Franklin Lakes, NJ) (see **Fig. S2** for gating strategies) and FlowJo software (BD, Version 10). Antibodies: CD45-eFluor 450 (Clone 30-F11), CD11c-PE-Cy7 (Clone N418), MHC II (I-A/I-E)-APC-eFlour780 (Clone M5/114.15.2), CD103-FITC (Clone 2E7), CD274 (B7-H1, PD-L1)-PE-Cy7 (Clone MIH5), CD3-FITC (Clone 17A2), CD8-eFluor450 (Clone 53-6.7), CD107a (LAMP-1)-PE (Clone 1D4B), IFN-γ-PE (Clone XMG1.2), Granzyme B-FITC (Clone NGZB), γδ TCR-PE-Cy5 (Clone GL3), and FoxP3-PE (Clone NRRF-30) all from eBioscience (Waltham, MA); CD4-PerCP-Cy5.5 (Clone BM9), CD69-PE (Clone H1.2F3), F4/80-PerCP-Cy5.5 (Clone BM9), MHC I (H-2)-PE (Clone M1/42), CD326 (EpCAM)-APC (Clone G8.8), and CD86-BV510 (Clone GL-1) all from BioLegend; CD45-APC (Clone 30-F11), Ly6G-FITC (Clone 1A8), and CD11b-PE (Clone M1/70) from BD Pharmingen; EpCAM-APC (Clone CI:A3-1, AbD Serotec [Hercules, CA]) and CCR2-flourescein (Clone 475301, R&D Systems). Prior to Granzyme B staining, cells were incubated for 2 h with Brefeldin A (Sigma-Aldrich, #B6542).

### Cytokine array

Tumors (8-10 mm in diameter) were isolated from MMTV-PyMT;*Ccr2*^+/+^ and MMTV-PyMT;*Ccr2*^−/−^ mice and immediately snap-frozen in liquid nitrogen. They were then homogenized in PBS with the addition of protease inhibitors (Promega, Madison, WI). Proteome Profiler Mouse Cytokine Array Kit, Panel A (R&D Systems) was used according to the manufacturer’s instructions. Films were scanned, and spots were analyzed using ImageJ software.

### MTS assay

Proliferation of cancer cells isolated from MMTV-PyMT;*Ccr2*^+/+^ and MMTV-PyMT;*Ccr2*^−/−^ mice was measured using the CellTiter 96® AQueous One Solution Cell Proliferation Assay (MTS; Promega) according to the manufacturer’s instructions. In this assay, the MTS tetrazolium compound (Owen’s reagent) was bioreduced by metabolically active cells into a colored formazan product. The quantity of formazan product was measured in a 96-well plate by absorbance at 490 nm.

### RNA purification and real-time quantitative PCR (RT-qPCR) analysis

Total RNA from cancer cells (cultured 48 hours in RPMI supplemented with 10% FBS) or primary tumors was extracted using an RNAeasy kit (Qiagen, Germantown, MD), following the manufacturer’s instructions. RNA was quantified using a Nanodrop 2000 spectrophotometer (Thermo Fisher Scientific, Waltham, MA). Reverse transcription was performed with an ImProm-II Reverse Transcription System (Promega) for the cancer cells, or the RevertAid First Strand cDNA Synthesis system (Thermo, K1622) for the primary tumors, with the following cycling conditions: 5 min at 70°C, 10 min at 4°C, 5 min at 25°C, 60 min at 42°C, and 15 sec at 70°C using 1 μg of total RNA extract for 20 μL of final volume. RT-qPCR analysis was performed using a TaqMan gene expression assay (Thermo Fisher Scientific) with the following specific primers: *Ccr2*: Mm00438270_m1; *Pd-l1:* Mm00452054_m1; *Cd8a*: Mm01182107_g1; *Ifn-g:* mm01168134_m1; *Pd-1*: mm01285676_m1; *Gzmb*: mm00502528_m1; *Ctla4*: mm00486849_m1; *Prf1*: mm00812512_m1; *Clec9a*: mm00554956_m1; *Cxcl9*: mm00434946_m1; *Cxcl10*: mm00445235_m1; *Cxcr3*: mm00438259_m1; *Cxcr4*: Mm01996749_s1; *Cd11c*: mm00498698_m1; *Cd40*: mm00441891_m1; *Cd80*: mm00711660_m1; *Cd86*: mm00444543_m1; *Ccl5*: mm01302427_m1; *Il-12b*: mm01288989_m1; *Tim3*: mm00454540_m1; *Csf2*: mm01290062_m1; *Flt3l*: mm00442801_m1; *Flt3*: mm00439016_m1; *Ccl2*: mm00441242_m1; *Csf3*: mm00438334_m1; *Csf1*: mm00432686_m1 and *Tbp*: mm01277042_m1. For cancer cells, RT-qPCR was performed on three independently isolated cancer cell populations and in triplicate for each sample. To determine the effects of IFN-γ on *Pd-l1* expression, cancer cells were cultured with indicated concentration of IFN-γ (R&D systems, catalogue #485-MI-100) for 48 h before isolation of RNA. For tumor samples, RT-qPCR was performed on at least five primary tumors from different mice and in triplicate for each sample. Relative quantitation was performed with the 2^(−ΔΔCT)^ method using *β-Actin* or *Tbp* expression for normalization and MMTV-PyMT;*Ccr2*^+/+^ cancer cells or tumor samples as a reference sample.

### RNA *in situ* hybridization

RNA *in situ* hybridization (ISH) was performed on PFA-fixed paraffin-embedded tissue sections using an RNAscope Chromogenic 2.0 Detection Kit (Advanced Cell Diagnostics [ACD], Inc., Newark, CA). A *Ccr2* probe was custom-designed by ACD, Inc. (Catalog number 436261-C2) and used according to the manufacturer’s instructions. We confirmed that no signal was found with the probe using *Ccr2^−/−^* tissues. RNA fluorescent ISH (FISH) for *Ccr2, Krt8, Cd8a,* and *Ifn-g* was performed on fresh frozen tissue sections using an RNAscope Fluorescent Multiplex Assay Kit (Advanced Cell Diagnostics [ACD], Inc., Newark, CA) with the following probes (by ACD): *Ccr2*: 436261-C2; *Krt18*: 424531-C1; *Cd8a*: 401681-C1; and *Ifn-g*: 311391-C3) and used according to the manufacturer’s instructions. For *Ccr2* RNA FISH, we confirmed that no signal was found with the probe using *Ccr2^−/−^* tissues.

### Hematoxylin and eosin staining

PFA-fixed paraffin-embedded tissue sections were deparaffinized and rehydrated following standard protocols, stained with Gill’s Hematoxylin (Thermo Fisher Scientific, REF6765008) for 3 min, washed in tap water, rinsed with Nu-Clear II (Thermo Fisher Scientific, REF6769009), and then rinsed again first with tap water and then with Bluing Reagent (Thermo Fisher Scientific, REF 6769001). Slides were then stained with eosin (Sigma, HT110180), dehydrated following standard protocols, air-dried, and mounted with Cytoseal 60 (Thermo Fisher Scientific, Catalog number 831016).

### Immunofluorescence staining

To stain for cytokeratin 5 and cytokeratin 8, paraffin sections were deparaffinized and rehydrated, and antigen retrieval was carried out by boiling slides in Tris EDTA buffer (10 mM Tris Base, 1 mM EDTA, pH 9.0) for 6 min in a pressure cooker. The slides were blocked with 1x blocking buffer (PBS containing 2.5% BSA and 5% goat serum) and Fc receptor blocker (Innovex Biosciences, Richmond, CA), before incubating with anti-cytokeratin 5-Alexa Fluor 488 conjugated antibody (ab193894; 1:200 dilution; Abcam) and anti-Cytokeratin 8-Alexa Fluor 405 conjugated antibody (ab210139; 1:200 dilution; Abcam) overnight at 4°C. Sections were counterstained with ToPro-3 (1:1000 dilution; Life Technologies).

For all other stainings, frozen tissue sections were incubated with 1x blocking buffer (5% goat serum, 2.5% BSA in PBS) and Fc receptor blocker (Innovex Biosciences). Sections were incubated with rabbit anti-CD3 polyclonal antibody (1:500 dilution; Abcam, Cambridge, MA), rat anti-MHC class I monoclonal antibody (Clone ER-HR52, 1:100 dilution; Abcam), rat anti-PyMT antibody (Ab15085, 1:200; Abcam), rabbit anti-cleaved caspase-3 (Asp175) (D3E9) (#9579; 1:200 dilution; Cell Signaling Technology), anti-CD8a-Alexa Fluor 488 (Clone 5.3-6.7), anti-CD11c-Alexa Fluor 488 (Clone N418), or anti-CD326 (EpCAM)-APC (Clone G8.8) in 0.5 x blocking buffer overnight at 4°C. Secondary antibodies (not used for primary antibodies conjugated to Alexa-Fluor 488) anti-rabbit-Alexa568, or anti-rat-Alexa568 (1:150 dilution; Life Technologies) were used for detection, and sections were counterstained with DAPI (1:200 dilution; Life Technologies). Images were collected at 40x magnification using a Leica SP8 confocal or an AX10 microscope and an AxioCam HRc camera (Zeiss, Thornwood, NY) and were analyzed using Volocity software (Version 6.3.0, PerkinElmer, Waltham, MA).

### Immunohistochemistry and terminal deoxynucleotidyl transferase dUTP nick end labeling (TUNEL) staining

Sections were deparaffinized and rehydrated. For phospho-STAT3 and Ki67 staining, antigen retrieval was carried out by boiling slides in 10 mM sodium citrate buffer (pH 6.0) for 6 min in a pressure cooker, while for phospho-STAT1 staining, antigen retrieval was done in Tris EDTA buffer (10 mM Tris Base, 1 mM EDTA, pH 9.0). The slides were blocked with 3% hydrogen peroxide, 1x blocking buffer (PBS containing 2.5% BSA and 5% goat serum), Fc receptor blocker (Innovex Biosciences), and finally avidin/biotin blocking buffer (Vector Laboratories, Burlingame, CA, SP-2001). Slides were then incubated with primary antibodies overnight at 4°C in 0.5x blocking buffer: rabbit anti-Ki67/MK167 (1:1,000 dilution; Novus International, Saint Charles, MO, NB110-89717), rabbit anti-phospho-STAT1 (1:200 dilution; Cell Signaling Technology, Danvers, MA), or rabbit anti-phospho-STAT3 (1:500 dilution; Cell Signaling Technology). After incubating with secondary biotinylated goat anti-rabbit IgG antibody (1:500, Vector Laboratories, BA-1000) for 1 h at room temperature, slides were incubated with avidin-conjugated horseradish peroxidase (Vector Laboratories, PK-6100) for 30 min, and the signals were detected by a DAB (3,3’-Diaminobenzidine) substrate kit (Vector Laboratories, SK-4100). Lastly, sections were counter-stained with hematoxylin. TUNEL staining was performed on deparaffinized sections to detect late-stage apoptotic cells using the ApopTag peroxidase *in situ* apoptosis detection kit (Millipore, Burlington, MA), according to the manufacturer’s instructions. Images were scanned by an Aperio ScanScope® CS System (Leica Biosystems), and positive nuclei were counted using Aperio eSlide Capture Devices software (Leica Biosystems).

### ImmunoSEQ

Tumors from *Ccr2*^+/+^ and *Ccr2*^−/−^ transplants were isolated and immediately placed in liquid nitrogen for shipment. Amplification and sequencing were performed on the immunoSEQ platform (Adaptive Biotechnologies, Seattle, WA) using a multiplex PCR-based assay that exclusively targets rearranged T cell receptor genes. Sequencing data were analyzed using immunoSEQ Analyzer software. Shannon’s Entropy (H), a measure of the richness and uniformity of the frequency of the T cell receptor repertoire distribution, was defined as:

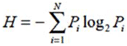

Where N equals the number of unique clones and P_i_ the frequency of clones. To account for variation in sequencing depth, entropy was normalized by its maximum value (H_max_):

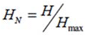

Clonality is defined as C = 1 - H_N_. (Sherwood et al., 2013). Sequencing files are available from the Adaptive Biotechnologies immunoSEQ web site.

### Transcriptional profiling and bioinformatics analysis

*Ccr2^+/+^* or *Ccr2^−/−^* cancer cells were isolated from tumors and sorted as EpCAM+CD45-CD31- live cells (using 7-AAD Viability Staining Solution, Thermo Fisher Scientific) four weeks after orthotopic transplantation into the mammary glands of wild-type C57BL/6 mice. Total cancer cell RNA was extracted using a TRIzol extraction protocol. cDNA libraries were prepared using the Ovation^®^ RNA-Seq System V2 (NuGEN Technologies, San Carlos, CA) according to the manufacturer’s protocol and sequenced using a NextSeq 500 instrument (Illumina, San Diego, CA) to obtain 388 million 76 bp single-end reads. Reads were mapped to the reference mm9 mouse genome using Spliced Transcripts Alignment to a Reference (STAR) (Dobin et al., 2013) and were determined using HTSeq (Anders et al., 2015). Differential gene expression analysis was performed using DESeq2 (Love et al., 2014) with a false discovery rate cutoff of 5%. Hierarchical clustering was performed by sample and gene using normalized and log2-transformed gene expression values of differentially expressed genes. All of the analyses described above were performed on the Bioinformatics Shared Resource Galaxy server at CSHL. Gene set enrichment analysis (Subramanian et al., 2005) was performed on a pre-ranked gene list sorted by log2FoldChange value against v5.1 Hallmark gene sets using default parameters. Gene ontology term enrichment analysis was performed using GOrilla PANTHER (Eden et al., 2009).

### Bone marrow-derived dendritic cell (BMDC) isolation, culture, and activation

BMDCs were isolated and cultured as previously described (Inaba et al., 1992). Briefly, bone marrow was obtained from female C57BL/6 mice by flushing the femur and tibia with 2 ml of HBSS using a 1 mL insulin syringe with a 29G × ½ needle. The cells were washed with HBSS twice by centrifugation at 250 x g for 8 min and then suspended in BMDC culture medium (RPMI-1640 medium [Thermo Fisher Scientific] containing 10% FBS, 20 ng/ml recombinant mouse GM-CSF [BioLegend], 50 units/ml penicillin, and 50 μg/ml streptomycin). The cells were then plated in 10 cm culture plates (2×10^6^ cells per petri dish) and kept at 37°C in a 5% CO_2_ incubator for three days, then split at day 4 into two plates. For activation of BMDCs with cancer cell conditioned medium, the non-attached BMDCs were harvested at day 6 by gently pipetting the cultures with medium. This BMDC population was then rinsed three times in HBSS and suspended in a) culture medium, b) conditioned medium from MMTV-PyMT;*Ccr2*^−/−^ or MMTV-PyMT;*Ccr2*^+/+^ cancer cells, or c) conditioned medium from MMTV-PyMT;*Ccr2*^+/+^ cancer cells cultured (either with or without CCR2 antagonist (RS504393, 10 μM, Sigma-Aldrich). After incubation with conditioned medium for 24 h, non-adherent BMDCs were harvested for flow cytometry.

### Chromium release assay

Cytotoxic CD8+ T cells were isolated from spleens of OT-I transgenic mice by magnetic labeling and separation (Miltenyi Biotec, Cambridge, MA). Briefly, the spleen was forced through a 100 μm cell strainer (BD Biosciences), and flow-through cells were collected. After lysing red blood cells, the splenocytes were washed and incubated with CD8a+ T Cell Biotin-Antibody Cocktail, mouse (Miltenyi Biotec) at 4°C for 5 min, followed by incubating with anti-Biotin microbeads (Miltenyi Biotec) at 4°C for 10 min. The cell suspension was applied to an LS column (Miltenyi Biotec) in a magnetic field, and the flow-through (negatively selected) CD8+ T cells were collected. The isolated CD8+ T cells were cultured on plates coated with anti-mouse-CD3 (BioLegend) in RPMI-1640 containing 10% FBS, 50 μM of β-mercaptoethanol, 20 ng/ml mouse IL-2, 2 ng/ml mouse IL-7, and 1% penicillin/streptomycin for 6-9 days. Primary cancer cells were isolated as described above from MMTV-PyMT-chOVA;*Ccr2*^+/+^ or MMTV-PyMT-chOVA;*Ccr2*^−/−^ mice and cultured overnight with or without CCR2 antagonist (10 μM, Sigma RS504393) and OVA peptides (1 μg/ml) in DMEM/F12 medium with 10% FBS. The E0771 cancer cell line was infected with lentivirus expressing luciferase and OVA SIINFEKL peptide fused to the C-terminus of eGFP. Transduced cells were sorted by flow cytometry for eGFP-positive cells. For chromium (^51^Cr) release assay, cancer cells (1,000 per well) were incubated with 50 μl of ^51^Cr solution (activity 1 mCi/ml, PerkinElmer) plus 50 μl of RPMI medium with 10% FBS for 2 h at 37°C. After rinsing the cells three times with RPMI plus 10% FBS, 50 μl of cancer cells (1,000) were plated into each well of a 96-well, V-bottom plate, followed by the addition of 50 μl of CD8+ T cells at a ratio of 10:1. After 4 h of incubation at 37°C, 50 μl of supernatant from each well was transferred to miniature vials, and radioactivity was determined by a Beckman scintillation counter (PerkinElmer). The percentage of specific lysis was calculated using the standard formula [(experimental - spontaneous release)/(total - spontaneous release) x 100] and expressed as the mean of triplicate samples.

### Western blot

Snap-frozen tumor samples were homogenized in radio immunoprecipitation assay buffer with cOmplete^TM^ Protease Inhibitor Cocktail Tablets (Roche) and phosphatase inhibitor cocktail (Thermo Fisher Scientific), and lysis was performed on ice for 30 min. Cell lysates were centrifuged for 15 min at 12,000 x g at 4°C. After centrifugation, the supernatant was mixed with 4x loading buffer plus 2.5% of β-mercaptoethanol and loaded on 8% SDS/PAGE gels. Protein bands were transferred to PVDF membranes (Bio-RAD) at 90 V for 2 h in a Bio-Rad Mini Trans-Blot system. The membrane was incubated sequentially with different primary antibodies: anti-p65, anti-p-p65, anti-p-STAT1, anti-p-STAT3, and anti-STAT3 (all from Cell Signaling Technology), and anti-STAT1 and anti-beta-Actin (from Santa Cruz Biotechnology, Dallas, Tx). Secondary antibodies were from LI-COR, (Lincoln, NE). Protein detection was performed using the Odyssey imaging system (LI-COR).

### ELISA

Conditioned medium from *Ccr2^+/+^* and *Ccr2^−/−^* cancer cells was collected, and GM-CSF concentration was evaluated by ELISA, following the manufacturer’s instructions (R&D Systems). Briefly, 50 µl of cytokine standard or conditioned medium was added to the wells of the pre-coated ELISA plate and incubated at room temperature for 2 h. Plates were washed five times with wash buffer and incubated with a horseradish peroxidase-linked polyclonal antibody specific to mouse GM-CSF for 2 h, followed by incubation with substrate solution for 30 min. After adding 100 μL of stop solution, the optical density of each well was measured at 450 nm using a microplate reader. The concentrations of GM-CSF in the conditioned medium were calculated according to the standard curve.

### Whole mount carmine staining of mammary glands

Mammary glands were isolated and spread out on glass slides. Then, the mammary glands were fixed in Carnoy’s fixative (60% ethanol, 30% chloroform, and 10% glacial acetic acid) for 2–4 hours at room temperature, followed by 15-minute washes first in 70% and then in 50% ethanol, with a final rinse in double-distilled water. The glands were then stained with carmine red stain (2.5 g of aluminum potassium sulfate, 1 g of carmine, and dH_2_O to a final volume of 500 mL, boiled for 25 min) for 16 hours or until the fat pad was a uniform pink color. After staining, the mammary glands were de-stained in 1% solution of 1N HCl in 70% ethanol. De-stained glands were rinsed twice for 15 min in 70% ethanol followed by 15-minute washes in first in 90% and then in 100% ethanol. The mammary glands were then placed in xylene for two hours for clearing and mounted.

### Statistical analysis

All statistical analyses were performed using GraphPad Prism Version 6 software. Data were analyzed using Kaplan-Meier survival analysis, Student’s t-tests, ANOVA, two-way ANOVA, chi-Square test, and log-rank (Mantel-Cox) test, as indicated in the figure legends, with an alpha of 0.05. The number of sampled units, N, is indicated in the figure legends.

## ACKNOWLEDGEMENTS

We thank Laura Maiorino for drawing the included illustrations. This work was supported by funds from the CSHL Cancer Center (P30-CA045508) and the Department of Defense (W81XWH-14-1-0078) to M.E.; from the Charles and Marie Robertson Foundation to X.Y.H.; from the Lustgarten Foundation, the National Cancer Institute, and the Cedar Hill Foundation to D.T.F.; from the Rita Allen Foundation and the V Foundation for Cancer Research to C.O.d.S.; a Starr Centennial Scholarship from the Watson School of Biological Sciences to E.B; and R.Y is a George A. and Marjorie H. Anderson Fellow. Support was also provided by the State of New York (Contract #C150158), and the opinions, results, findings and/or interpretations of data contained therein do not necessarily represent the opinions, interpretations or policy of the State.

## AUTHOR CONTRIBUTIONS

M.R.F., X.Y.H., and M.E. designed the experiments. M.R.F. and X.Y.H. performed the *in vitro* experiments; M.R.F., C.O.d.S., and E.B. isolated RNA from cancer cells and performed the RNA-seq analysis. M.R.F. and X.Y.H. performed the animal experiments; A.S.A., M.R.F., A.P., and X.Y.H. performed flow cytometry; J.E.W. analyzed the histological sections; and M.R.F., X.Y.H, and A.E. performed immunohistochemistry and *in situ* hybridizations. X.Y.H., A.P., R.Y., and M.R.F. designed and performed the chromium release experiments; and A.E., M.R.F., and X.Y.H. performed qPCR. L.V.A. and D.T.F. provided experimental advice. M.R.F., X.Y.H., and M.E. wrote the manuscript.

## Abbreviations

BMDC: bone marrow-derived dendritic cells
CCL2: C-C chemokine ligand 2
CCR2: C-C chemokine receptor type 2
chOVA: mCherry-ovalbumin
CTL: cytotoxic T lymphocyte
DC: dendritic cell
dLN: draining lymph node
FISH: Fluorescence *in situ* hybridization
G-CSF: granulocyte colony-stimulating factor
GM-CSF: granulocyte-macrophage colony-stimulating factor
IACUC: Institutional Animal Care and Use Committee
ISH: *in situ* hybridization
IFN-γ: interferon gamma
LAMP-1: lysosomal-associated membrane protein 1
MDSC: myeloid-derived suppressor cell
MHC: major histocompatibility complex
MMTV: mouse mammary tumor virus
PD-L1: programmed cell death ligand 1
PFA: paraformaldehyde
PyMT: polyoma middle T
RPKM: reads per kilobase million
RPMI: Roswell Park Memorial Institute
TCGA: The Cancer Genome Atlas
TCR: T cell receptor
TUNEL: terminal deoxynucleotidyl transferase dUTP nick end labelling

## REFERENCES

Ahn, R., V. Sabourin, A.M. Bolt, S. Hebert, S. Totten, N. De Jay, M.C. Festa, Y.K. Young, Y.K. Im, T. Pawson, A.E. Koromilas, W.J. Muller, K.K. Mann, C.L. Kleinman, and J. Ursini-Siegel. 2017. The Shc1 adaptor simultaneously balances Stat1 and Stat3 activity to promote breast cancer immune suppression. Nat Commun 8:14638.

Anders, S., P.T. Pyl, and W. Huber. 2015. HTSeq--a Python framework to work with high-throughput sequencing data. Bioinformatics 31:166–169.

Bayne, L.J., G.L. Beatty, N. Jhala, C.E. Clark, A.D. Rhim, B.Z. Stanger, and R.H. Vonderheide. 2012. Tumor-derived granulocyte-macrophage colony-stimulating factor regulates myeloid inflammation and T cell immunity in pancreatic cancer. Cancer Cell 21:822–835.

Bonapace, L., M.M. Coissieux, J. Wyckoff, K.D. Mertz, Z. Varga, T. Junt, and M. Bentires-Alj. 2014. Cessation of CCL2 inhibition accelerates breast cancer metastasis by promoting angiogenesis. Nature 515:130–133.

Boring, L., J. Gosling, S.W. Chensue, S.L. Kunkel, R.V. Farese, Jr., H.E. Broxmeyer, and I.F. Charo. 1997. Impaired monocyte migration and reduced type 1 (Th1) cytokine responses in C-C chemokine receptor 2 knockout mice. J Clin Invest 100:2552–2561.

Broz, M.L., M. Binnewies, B. Boldajipour, A.E. Nelson, J.L. Pollack, D.J. Erle, A. Barczak, M.D. Rosenblum, A. Daud, D.L. Barber, S. Amigorena, L.J. Van’t Veer, A.I. Sperling, D.M. Wolf, and M.F. Krummel. 2014. Dissecting the tumor myeloid compartment reveals rare activating antigen-presenting cells critical for T cell immunity. Cancer Cell 26:638–652.

Chen, J., C.C. Jiang, L. Jin, and X.D. Zhang. 2016a. Regulation of PD-L1: a novel role of pro-survival signalling in cancer. Ann Oncol 27:409–416.

Chen, W., Q. Gao, S. Han, F. Pan, and W. Fan. 2015. The CCL2/CCR2 axis enhances IL-6-induced epithelial-mesenchymal transition by cooperatively activating STAT3-Twist signaling. Tumour Biol 36:973–981.

Chen, X., Y. Wang, D. Nelson, S. Tian, E. Mulvey, B. Patel, I. Conti, J. Jaen, and B.J. Rollins. 2016b. CCL2/CCR2 Regulates the Tumor Microenvironment in HER-2/neu-Driven Mammary Carcinomas in Mice. PLoS One 11:e0165595.

de Mingo Pulido, A., A. Gardner, S. Hiebler, H. Soliman, H.S. Rugo, M.F. Krummel, L.M. Coussens, and B. Ruffell. 2018. TIM-3 Regulates CD103(+) Dendritic Cell Function and Response to Chemotherapy in Breast Cancer. Cancer Cell 33:60–74 e66.

Deshmane, S.L., S. Kremlev, S. Amini, and B.E. Sawaya. 2009. Monocyte chemoattractant protein-1 (MCP-1): an overview. J Interferon Cytokine Res 29:313–326.

Dobin, A., C.A. Davis, F. Schlesinger, J. Drenkow, C. Zaleski, S. Jha, P. Batut, M. Chaisson, and T.R. Gingeras. 2013. STAR: ultrafast universal RNA-seq aligner. Bioinformatics 29:15–21.

Dunn, G.P., A.T. Bruce, H. Ikeda, L.J. Old, and R.D. Schreiber. 2002. Cancer immunoediting: from immunosurveillance to tumor escape. Nat Immunol 3:991–998.

Dunn, G.P., L.J. Old, and R.D. Schreiber. 2004. The three Es of cancer immunoediting. Annu Rev Immunol 22:329–360.

Eden, E., R. Navon, I. Steinfeld, D. Lipson, and Z. Yakhini. 2009. GOrilla: a tool for discovery and visualization of enriched GO terms in ranked gene lists. BMC Bioinformatics 10:48.

Engelhardt, J.J., B. Boldajipour, P. Beemiller, P. Pandurangi, C. Sorensen, Z. Werb, M. Egeblad, and M.F. Krummel. 2012. Marginating dendritic cells of the tumor microenvironment cross-present tumor antigens and stably engage tumor-specific T cells. Cancer Cell 21:402–417.

Fang, W.B., I. Jokar, A. Zou, D. Lambert, P. Dendukuri, and N. Cheng. 2012. CCL2/CCR2 chemokine signaling coordinates survival and motility of breast cancer cells through Smad3 protein- and p42/44 mitogen-activated protein kinase (MAPK)-dependent mechanisms. J Biol Chem 287:36593–36608.

Freeman, G.J., A.J. Long, Y. Iwai, K. Bourque, T. Chernova, H. Nishimura, L.J. Fitz, N. Malenkovich, T. Okazaki, M.C. Byrne, H.F. Horton, L. Fouser, L. Carter, V. Ling, M.R. Bowman, B.M. Carreno, M. Collins, C.R. Wood, and T. Honjo. 2000. Engagement of the PD-1 immunoinhibitory receptor by a novel B7 family member leads to negative regulation of lymphocyte activation. J Exp Med 192:1027–1034.

Garrido, F., I. Romero, N. Aptsiauri, and A.M. Garcia-Lora. 2016. Generation of MHC class I diversity in primary tumors and selection of the malignant phenotype. Int J Cancer 138:271–280.

Greter, M., J. Helft, A. Chow, D. Hashimoto, A. Mortha, J. Agudo-Cantero, M. Bogunovic, E.L. Gautier, J. Miller, M. Leboeuf, G. Lu, C. Aloman, B.D. Brown, J.W. Pollard, H. Xiong, G.J. Randolph, J.E. Chipuk, P.S. Frenette, and M. Merad. 2012. GM-CSF controls nonlymphoid tissue dendritic cell homeostasis but is dispensable for the differentiation of inflammatory dendritic cells. Immunity 36:1031–1046.

Guy, C.T., R.D. Cardiff, and W.J. Muller. 1992. Induction of mammary tumors by expression of polyomavirus middle T oncogene: a transgenic mouse model for metastatic disease. Mol Cell Biol 12:954–961.

Hildner, K., B.T. Edelson, W.E. Purtha, M. Diamond, H. Matsushita, M. Kohyama, B. Calderon, B.U. Schraml, E.R. Unanue, M.S. Diamond, R.D. Schreiber, T.L. Murphy, and K.M. Murphy. 2008. Batf3 deficiency reveals a critical role for CD8alpha+ dendritic cells in cytotoxic T cell immunity. Science 322:1097–1100.

Hu, Q., M. Myers, W. Fang, M. Yao, G. Brummer, J. Hawj, C. Smart, C. Berkland, and N. Cheng. 2019. Role of ALDH1A1 and HTRA2 expression in CCL2/CCR2-mediated breast cancer cell growth and invasion. Biol Open 8: bio040873

Igney, F.H., and P.H. Krammer. 2002. Immune escape of tumors: apoptosis resistance and tumor counterattack. J Leukoc Biol 71:907–920.

Inaba, K., M. Inaba, N. Romani, H. Aya, M. Deguchi, S. Ikehara, S. Muramatsu, and R.M. Steinman. 1992. Generation of large numbers of dendritic cells from mouse bone marrow cultures supplemented with granulocyte/macrophage colony-stimulating factor. J Exp Med 176:1693–1702.

Izumi, K., L.Y. Fang, A. Mizokami, M. Namiki, L. Li, W.J. Lin, and C. Chang. 2013. Targeting the androgen receptor with siRNA promotes prostate cancer metastasis through enhanced macrophage recruitment via CCL2/CCR2-induced STAT3 activation. EMBO Mol Med 5:1383–1401.

Jones, L.M., M.L. Broz, J.J. Ranger, J. Ozcelik, R. Ahn, D. Zuo, J. Ursini-Siegel, M.T. Hallett, M. Krummel, and W.J. Muller. 2016. STAT3 Establishes an Immunosuppressive Microenvironment during the Early Stages of Breast Carcinogenesis to Promote Tumor Growth and Metastasis. Cancer Res 76:1416–1428.

Labovsky, V., L.M. Martinez, K.M. Davies, M. de Lujan Calcagno, H. Garcia-Rivello, A. Wernicke, L. Feldman, A. Matas, M.B. Giorello, F.R. Borzone, H. Choi, S.C. Howard, and N.A. Chasseing. 2017. Prognostic significance of TRAIL-R3 and CCR-2 expression in tumor epithelial cells of patients with early breast cancer. BMC Cancer 17:280.

Lebrecht, A., C. Grimm, T. Lantzsch, E. Ludwig, L. Hefler, E. Ulbrich, and H. Koelbl. 2004. Monocyte chemoattractant protein-1 serum levels in patients with breast cancer. Tumour Biol 25:14–17.

Lebrecht, A., L. Hefler, C. Tempfer, and H. Koelbl. 2001. Serum cytokine concentrations in patients with cervical cancer: interleukin-4, interferon-gamma, and monocyte chemoattractant protein-1. Gynecol Oncol 83:170–171.

Li, M., D.A. Knight, A.S. L, M.J. Smyth, and T.J. Stewart. 2013. A role for CCL2 in both tumor progression and immunosurveillance. Oncoimmunology 2:e25474.

Lim, S.Y., A.E. Yuzhalin, A.N. Gordon-Weeks, and R.J. Muschel. 2016. Targeting the CCL2-CCR2 signaling axis in cancer metastasis. Oncotarget 7: 28697–28710.

Lin, E.Y., J.G. Jones, P. Li, L. Zhu, K.D. Whitney, W.J. Muller, and J.W. Pollard. 2003. Progression to malignancy in the polyoma middle T oncoprotein mouse breast cancer model provides a reliable model for human diseases. Am J Pathol 163:2113–2126.

Love, M.I., W. Huber, and S. Anders. 2014. Moderated estimation of fold change and dispersion for RNA-seq data with DESeq2. Genome Biol 15:550.

Lu, Y., Z. Cai, D.L. Galson, G. Xiao, Y. Liu, D.E. George, M.F. Melhem, Z. Yao, and J. Zhang. 2006. Monocyte chemotactic protein-1 (MCP-1) acts as a paracrine and autocrine factor for prostate cancer growth and invasion. Prostate 66:1311–1318.

Mayer, C.T., P. Ghorbani, A. Nandan, M. Dudek, C. Arnold-Schrauf, C. Hesse, L. Berod, P. Stuve, F. Puttur, M. Merad, and T. Sparwasser. 2014. Selective and efficient generation of functional Batf3-dependent CD103+ dendritic cells from mouse bone marrow. Blood 124:3081–3091.

Mehta, H.M., M. Malandra, and S.J. Corey. 2015. G-CSF and GM-CSF in Neutropenia. J Immunol 195:1341–1349.

Meyer, M.A., J.M. Baer, B.L. Knolhoff, T.M. Nywening, R.Z. Panni, X. Su, K.N. Weilbaecher, W.G. Hawkins, C. Ma, R.C. Fields, D.C. Linehan, G.A. Challen, R. Faccio, R.L. Aft, and D.G. DeNardo. 2018. Breast and pancreatic cancer interrupt IRF8-dependent dendritic cell development to overcome immune surveillance. Nat Commun 9:1250.

Mitchell, L.A., R.J. Hansen, A.J. Beaupre, D.L. Gustafson, and S.W. Dow. 2013. Optimized dosing of a CCR2 antagonist for amplification of vaccine immunity. Int Immunopharmacol 15:357–363.

Muller, A., B. Homey, H. Soto, N. Ge, D. Catron, M.E. Buchanan, T. McClanahan, E. Murphy, W. Yuan, S.N. Wagner, J.L. Barrera, A. Mohar, E. Verastegui, and A. Zlotnik. 2001. Involvement of chemokine receptors in breast cancer metastasis. Nature 410:50–56.

Nakasone, E.S., H.A. Askautrud, T. Kees, J.H. Park, V. Plaks, A.J. Ewald, M. Fein, M.G. Rasch, Y.X. Tan, J. Qiu, J. Park, P. Sinha, M.J. Bissell, E. Frengen, Z. Werb, and M. Egeblad. 2012. Imaging tumor-stroma interactions during chemotherapy reveals contributions of the microenvironment to resistance. Cancer Cell 21:488–503.

Nielsen, B.S., L.R. Lund, I.J. Christensen, M. Johnsen, P.A. Usher, L. Wulf-Andersen, T.L. Frandsen, K. Dano, and H.J. Gundersen. 2001. A precise and efficient stereological method for determining murine lung metastasis volumes. Am J Pathol 158:1997–2003.

Nywening, T.M., A. Wang-Gillam, D.E. Sanford, B.A. Belt, R.Z. Panni, B.M. Cusworth, A.T. Toriola, R.K. Nieman, L.A. Worley, M. Yano, K.J. Fowler, A.C. Lockhart, R. Suresh, B.R. Tan, K.H. Lim, R.C. Fields, S.M. Strasberg, W.G. Hawkins, D.G. DeNardo, S.P. Goedegebuure, and D.C. Linehan. 2016. Targeting tumour-associated macrophages with CCR2 inhibition in combination with FOLFIRINOX in patients with borderline resectable and locally advanced pancreatic cancer: a single-centre, open-label, dose-finding, non-randomised, phase 1b trial. Lancet Oncol 17:651–662.

Pylayeva-Gupta, Y., K.E. Lee, C.H. Hajdu, G. Miller, and D. Bar-Sagi. 2012. Oncogenic Kras-induced GM-CSF production promotes the development of pancreatic neoplasia. Cancer Cell 21:836–847.

Qian, B., Y. Deng, J.H. Im, R.J. Muschel, Y. Zou, J. Li, R.A. Lang, and J.W. Pollard. 2009. A distinct macrophage population mediates metastatic breast cancer cell extravasation, establishment and growth. PLoS One 4:e6562.

Qian, B.Z., J. Li, H. Zhang, T. Kitamura, J. Zhang, L.R. Campion, E.A. Kaiser, L.A. Snyder, and J.W. Pollard. 2011. CCL2 recruits inflammatory monocytes to facilitate breast-tumour metastasis. Nature 475:222–225.

Roberts, E.W., M.L. Broz, M. Binnewies, M.B. Headley, A.E. Nelson, D.M. Wolf, T. Kaisho, D. Bogunovic, N. Bhardwaj, and M.F. Krummel. 2016. Critical Role for CD103(+)/CD141(+) Dendritic Cells Bearing CCR7 for Tumor Antigen Trafficking and Priming of T Cell Immunity in Melanoma. Cancer Cell 30:324–336.

Rodriguez-Frade, J.M., A.J. Vila-Coro, A.M. de Ana, J.P. Albar, A.C. Martinez, and M. Mellado. 1999. The chemokine monocyte chemoattractant protein-1 induces functional responses through dimerization of its receptor CCR2. Proc Natl Acad Sci U S A 96:3628–3633.

Ruffell, B., D. Chang-Strachan, V. Chan, A. Rosenbusch, C.M. Ho, N. Pryer, D. Daniel, E.S. Hwang, H.S. Rugo, and L.M. Coussens. 2014. Macrophage IL-10 blocks CD8+ T cell-dependent responses to chemotherapy by suppressing IL-12 expression in intratumoral dendritic cells. Cancer Cell 26:623–637.

Sherwood, A.M., R.O. Emerson, D. Scherer, N. Habermann, K. Buck, J. Staffa, C. Desmarais, N. Halama, D. Jaeger, P. Schirmacher, E. Herpel, M. Kloor, A. Ulrich, M. Schneider, C.M. Ulrich, and H. Robins. 2013. Tumor-infiltrating lymphocytes in colorectal tumors display a diversity of T cell receptor sequences that differ from the T cells in adjacent mucosal tissue. Cancer Immunol Immunother 62:1453–1461.

Soria, G., and A. Ben-Baruch. 2008. The inflammatory chemokines CCL2 and CCL5 in breast cancer. Cancer Lett 267:271–285.

Spranger, S., D. Dai, B. Horton, and T.F. Gajewski. 2017. Tumor-Residing Batf3 Dendritic Cells Are Required for Effector T Cell Trafficking and Adoptive T Cell Therapy. Cancer Cell 31:711–723 e714.

Subramanian, A., P. Tamayo, V.K. Mootha, S. Mukherjee, B.L. Ebert, M.A. Gillette, A. Paulovich, S.L. Pomeroy, T.R. Golub, E.S. Lander, and J.P. Mesirov. 2005. Gene set enrichment analysis: a knowledge-based approach for interpreting genome-wide expression profiles. Proc Natl Acad Sci U S A 102:15545–15550.

Winograd, R., K.T. Byrne, R.A. Evans, P.M. Odorizzi, A.R. Meyer, D.L. Bajor, C. Clendenin, B.Z. Stanger, E.E. Furth, E.J. Wherry, and R.H. Vonderheide. 2015. Induction of T-cell Immunity Overcomes Complete Resistance to PD-1 and CTLA-4 Blockade and Improves Survival in Pancreatic Carcinoma. Cancer Immunol Res 3:399–411.

Wolf, M.J., A. Hoos, J. Bauer, S. Boettcher, M. Knust, A. Weber, N. Simonavicius, C. Schneider, M. Lang, M. Sturzl, R.S. Croner, A. Konrad, M.G. Manz, H. Moch, A. Aguzzi, G. van Loo, M. Pasparakis, M. Prinz, L. Borsig, and M. Heikenwalder. 2012. Endothelial CCR2 Signaling Induced by Colon Carcinoma Cells Enables Extravasation via the JAK2-Stat5 and p38MAPK Pathway. Cancer Cell 22:91–105.

