## Supplemental figures for "Cancer cell CCR2 orchestrates suppression of the adaptive immune response"

### SUPPLEMENTAL FIGURE LEGENDS

#### Supplemental Figure 1: CCR2 expression in spontaneous MMTV-PyMT tumors and lungs

- **a.** Mammary ductal epithelial branching and invasion is similar in *Ccr2*$^{+/+}$ and *Ccr2*$^{-/-}$ mice, as indicated by carmine red staining of inguinal mammary glands at indicated age (left; Scale bar=4 mm) and immunofluorescence staining of basal and luminal cell markers (right; cytokeratin [CK] 5 and CK8, respectively; Scale bar=100 μm).
- **b.** Representative photomicrographs of H&E-stained tumors from MMTV-PyMT;*Ccr2*$^{+/+}$, MMTV-PyMT;*Ccr2*$^{+/-}$, and MMTV-PyMT;*Ccr2*$^{-/-}$ mice (Scale bar=1 mm).
- **c.** Histology score of solid area of primary tumors from MMTV-PyMT;*Ccr2*$^{+/+}$, MMTV-PyMT;*Ccr2*$^{+/-}$ and MMTV-PyMT;*Ccr2*$^{-/-}$ mice (mean +/- SEM, Student's t-test; n=8 in MMTV-PyMT;*Ccr2*$^{+/+}$, n=11 for MMTV-PyMT;*Ccr2*$^{+/-}$, and n=12 MMTV-PyMT;*Ccr2*$^{-/-}$).
- **d.** Representative RNA FISH on MMTV-PyMT;*Ccr2*$^{+/+}$ tumors (left) showing that *Ccr2* mRNA (green) is expressed by *Krt18* (red)-positive cancer cells. No *Ccr2* expression is detected in MMTV-PyMT;*Ccr2*$^{-/-}$ tumors (right; Scale bar=20 μm).
- **e.** Purity of cancer cells is greater than 90% after cell isolation, as determined by flow cytometry for EpCAM (black, a marker expressed on epithelial-derived MMTV-PyMT cells) compared to background (gray). Representative of three separate isolations.
- **f.** mRNA was collected from MMTV-PyMT;*Ccr2*$^{+/+}$, MMTV-PyMT;*Ccr2*$^{+/-}$, and MMTV-PyMT;*Ccr2*$^{-/-}$ tumor cells, and qPCR was performed for *Ccr2* and normalized to β-actin expression (mean +/- SEM, $p<0.05$, one-way ANOVA; n=3).

**g.** Tumors from *Ccr2*$^{+/-}$ cancer cells phenocopy those from *Ccr2*$^{-/-}$ cancer cells, regardless of whether the host is *Ccr2*$^{+/+}$ or *Ccr2*$^{-/-}$. Tumor burden was determined by weekly caliper measurement (mean +/- SEM, two-way ANOVA; n=8 for all conditions).

**h.** Metastatic burden is unchanged in MMTV-PyMT;*Ccr2*$^{+/+}$, MMTV-PyMT;*Ccr2*$^{+/-}$, and MMTV-PyMT;*Ccr2*$^{-/-}$ mice, as determined by quantification from H&E-stained lung sections, indicated as percentage of the area of the lung tissues (mean +/- SEM, ANOVA; n=24 MMTV-PyMT;*Ccr2*$^{+/+}$, 33 MMTV-PyMT;*Ccr2*$^{+/-}$, and 27 MMTV-PyMT;*Ccr2*$^{-/-}$ mice).

**i.** The number of metastatic foci is unchanged in MMTV-PyMT;*Ccr2*$^{+/+}$, MMTV-PyMT;*Ccr2*$^{+/-}$, and MMTV-PyMT;*Ccr2*$^{-/-}$ mice. Mice were classified as having either ≤ or > than 0.1 foci/mm$^2$ lung area (Chi-square test; n=24 MMTV-PyMT;*Ccr2*$^{+/+}$, 33 MMTV-PyMT;*Ccr2*$^{+/-}$, and 27 MMTV-PyMT;*Ccr2*$^{-/-}$ mice).

**j.** The average size of the metastatic foci is decreased in MMTV-PyMT;*Ccr2*$^{+/-}$ and MMTV-PyMT;*Ccr2*$^{-/-}$ mice compared to foci in MMTV-PyMT;*Ccr2*$^{+/+}$ mice. The number of mice with an average of large vs. small metastatic foci is indicated (Chi-square test; n=24 MMTV-PyMT;*Ccr2*$^{+/+}$, 33 MMTV-PyMT;*Ccr2*$^{+/-}$, and 27 MMTV-PyMT;*Ccr2*$^{-/-}$ mice).

**Supplemental Figure 2: Gating strategy for immune cells**

Gating strategy for immune cells. For flow cytometry analysis, single cells from tumors were gated on live CD45+ cells and were further analyzed using the indicated markers to characterize infiltration of tumor-associated macrophages (TAMs), DCs, T cells, granulocytes, monocytes, and MDSCs. SSC=side scatter.

**Supplemental Figure 3: T cell infiltration in tumors derived from *Ccr2*$^{+/+}$ or *Ccr2*$^{-/-}$ cancer cells**

**a.** There is no difference between *Ccr2*$^{+/+}$ and *Ccr2*$^{-/-}$ tumors during the growth-restricted phase in the percentage of CD3+ T cells among CD45+ leukocytes, as determined by flow cytometry for CD45+CD3+ cells (mean +/- SEM, Student's t-test; n=5).

**b.** FoxP3+CD4+ regulatory T cell infiltration is increased in *Ccr2*$^{+/+}$ tumors compared to *Ccr2*$^{-/-}$ tumors during the growth-restricted phase, as determined by flow cytometry gated on CD45+CD3+CD4+ cells (mean +/- SEM, Student's t-test; n=5).

**c.** Depletion of CD8+ T cells with anti-CD8a antibody significantly accelerated the growth of tumors from *Ccr2*$^{-/-}$ cancer cells (arrows indicate treatment with antibody; mean +/- SEM, two-way ANOVA; n=5 for all conditions).

**d.** Pharmacological inhibition of CCR2 increased OT-1 T cell cytotoxicity towards E0771 cancer cells expressing OVA peptide, as measured by chromium ($Cr^{51}$) release (mean +/- SEM, Student's t-test; each dot is a triplicate from the same experiment, and similar results were obtained in an independently performed experiment).

**e.** Immune suppression induced by cancer cell CCR2 signaling is confined to the local tumor microenvironment. *Ccr2*$^{+/+}$ cancer cells transplanted into one mammary gland did not alter the growth of tumors from *Ccr2*$^{-/-}$ cancer cells transplanted to the contralateral gland. Tumor burden was determined by weekly caliper measurement (mean +/- SEM, two-way ANOVA; n=10 for all conditions).

**Supplemental Figure 4: Myeloid cell infiltration in tumors derived from *Ccr2*$^{+/+}$ or *Ccr2*$^{-/-}$ cancer cells**

**a.** Tumors from transplanted *Ccr2*$^{-/-}$ cancer cells have elevated markers of T cell infiltration and activation, including *Cd8, Ifn-g,* and *Cxcl9* mRNA, compared to tumors from *Ccr2*$^{+/+}$ cancer cells. RT-qPCR was performed on tumors from transplants of *Ccr2*$^{+/+}$ and *Ccr2*$^{-/-}$ cancer cells isolated during the growth restricted phase (mean +/- SEM, Student's t-test; n=5–6 for *Ccr2*$^{+/+}$ tumors, and n=6–8 for *Ccr2*$^{-/-}$ tumors).

**b.** Representative RNA FISH for *Ifn-g* (green) and *Cd8a* (red) probes in the early phase (three weeks, upper panels) and the late, growth-restricted phase (5–6 weeks, bottom panels) of tumors from *Ccr2*$^{+/+}$ and *Ccr2*$^{-/-}$ transplanted cancer cells. Nuclei are counterstained with DAPI (blue; Scale bar=20 μm).

**c, d.** Quantification of RNA FISH shows more *Ifn-g*-expressing *Cd8+* T cells in tumors from *Ccr2*$^{-/-}$ cancer cells compared to *Ccr2*$^{+/+}$ cancer cells at both early (c) and late, growth-restricted (d) phases (mean +/- SEM; each dot represents the average of 3–8 random fields of view of one tumor; Student's t-test; n=5 tumors for early phase; n=7-8 tumors for late, growth-restricted phase).

**e.** Macrophage infiltration is unchanged between *Ccr2*$^{+/+}$ and *Ccr2*$^{-/-}$ tumors during the growth-restricted phase, as determined by flow cytometry gated on CD45+CD11b+MHC class II+ F4/80+ cells (mean +/- SEM, Student's t-test; n=5 and 4 for *Ccr2*$^{+/+}$ and *Ccr2*$^{-/-}$ tumors, respectively).

**f.** Percentage of CD11b+MHC class II- cells is reduced in *Ccr2*$^{-/-}$ tumors compared to *Ccr2*$^{+/+}$ tumors during the growth-restricted phase, as determined by flow cytometry gated on CD45+ cells (mean +/- SEM, Student's t-test; n=5 and 4 for *Ccr2*$^{+/+}$ and *Ccr2*$^{-/-}$ tumors, respectively).

**g.** Granulocytes and granulocytic MDSCs are decreased on *Ccr2*$^{-/-}$ tumors during the growth-restricted phase, as determined by flow cytometry gated on CD45+CD11b+CD11c-Ly6G+Ly6C+ cells (mean +/- SEM, Student's t-test; n=5).

**h.** Inflammatory monocytes and monocytic MDSCs are increased on *Ccr2*$^{-/-}$ tumors during the growth-restricted phase, as determined by flow cytometry gated on CD45+CD11b+CD11c-Ly6G-Ly6C+ cells (mean +/- SEM, Student's t-test; n=5).

**i.** CD103+ DCs are increased in MMTV-PyMT;*Ccr2*$^{-/-}$ tumors compared to MMTV-PyMT;*Ccr2*$^{+/+}$ tumors during the growth-restricted phase, as determined by flow cytometry gated on CD11c+MHCII+ cells within the CD45+ population (mean +/- SEM, Student's t-test; n=4 and 5 for MMTV-PyMT;*Ccr2*$^{+/+}$ and MMTV-PyMT;*Ccr2*$^{-/-}$ tumors, respectively).

**Supplemental Figure 5: Cytokine analysis in tumor lysate and conditioned medium**

**a.** Cytokine analysis from MMTV-PyMT;*Ccr2*$^{+/+}$ and MMTV-PyMT;*Ccr2*$^{-/-}$ tumors during growth-restricted phase, using the Proteome Profiler Mouse Cytokine Array Kit, Panel A (R&D Systems; mean +/- SEM, multiple Student's t-tests; n=4).

**b.** Cytokine analysis from supernatant of MMTV-PyMT;*Ccr2*$^{+/+}$ and MMTV-PyMT;*Ccr2*$^{-/-}$ cancer cells isolated during growth-restricted phase, using the Proteome Profiler Mouse Cytokine Array Kit, Panel A after serum starvation for 24 h (R&D Systems; mean +/- SEM, multiple Student's t-tests; n=4).

### Supplemental Figure 1

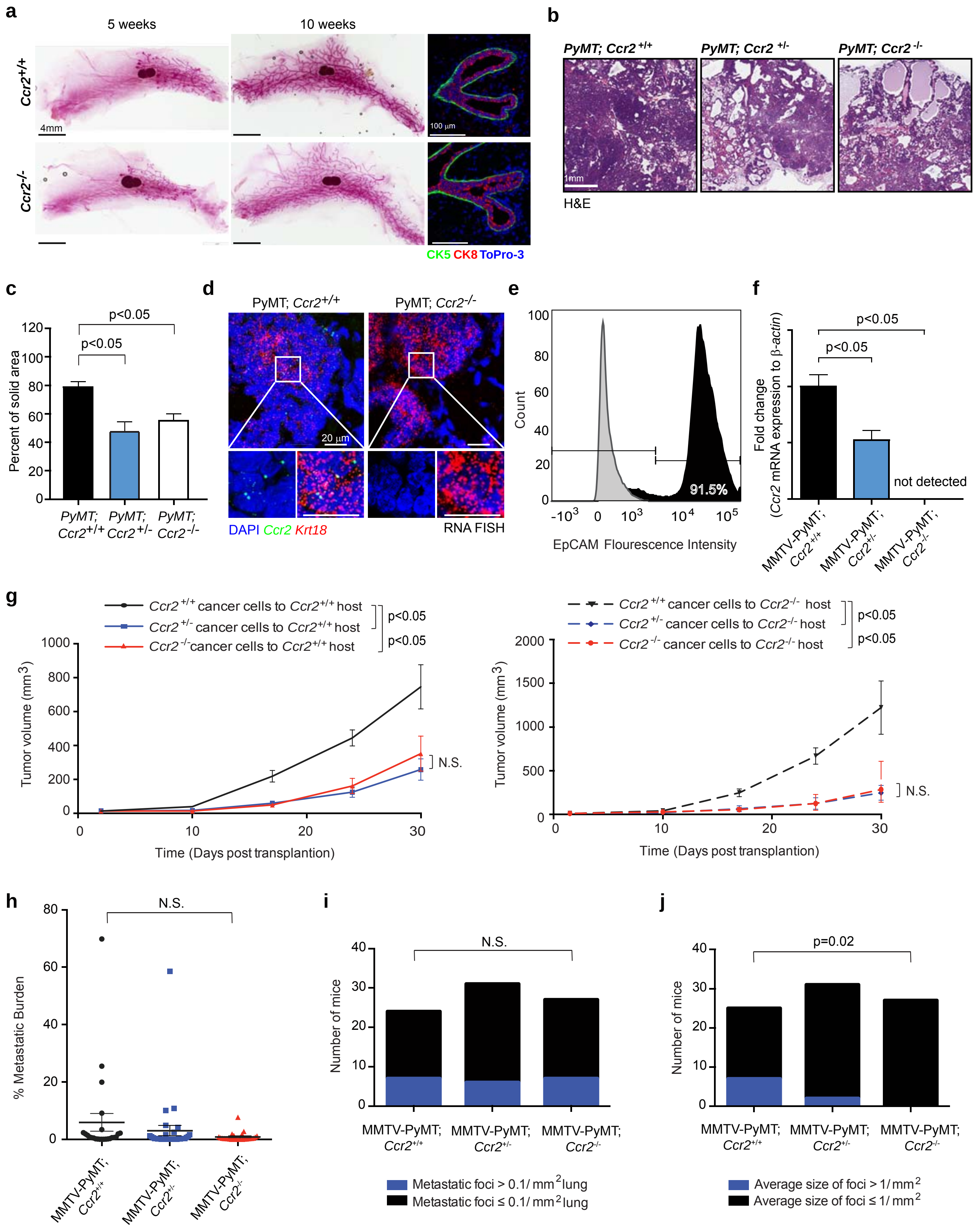

### Supplemental Figure 2

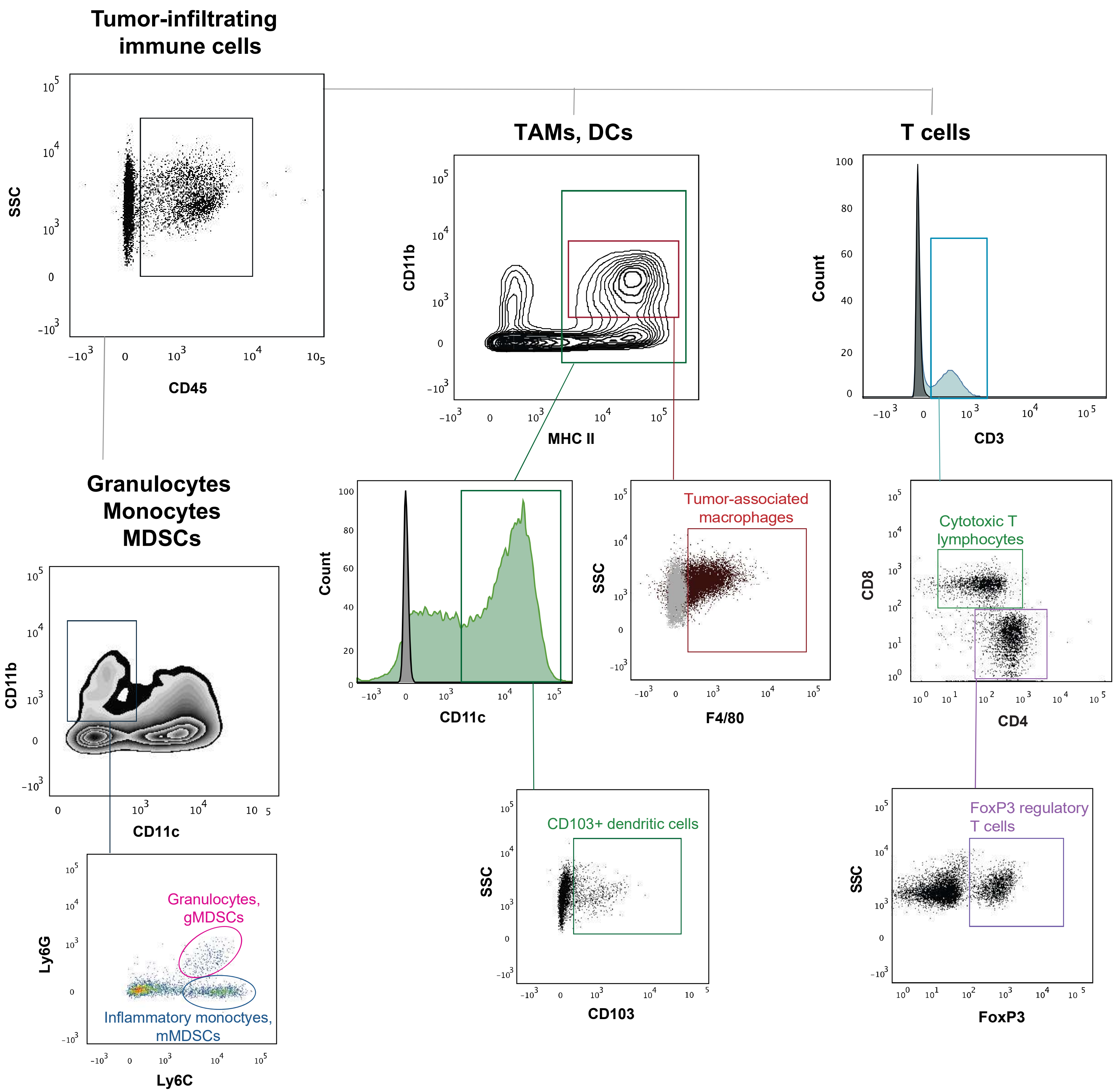

**Supplemental Figure 3**

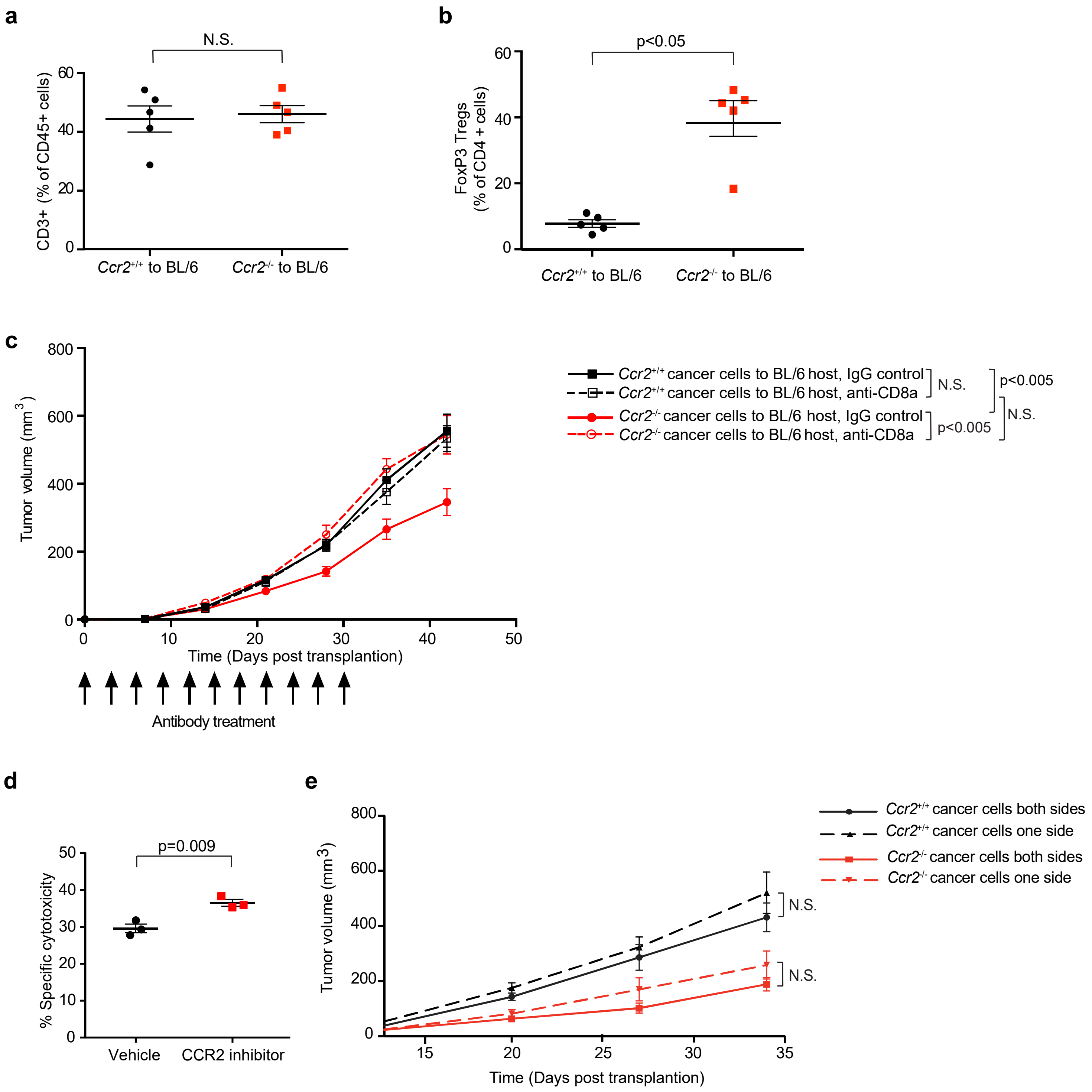

### Supplemental Figure 4

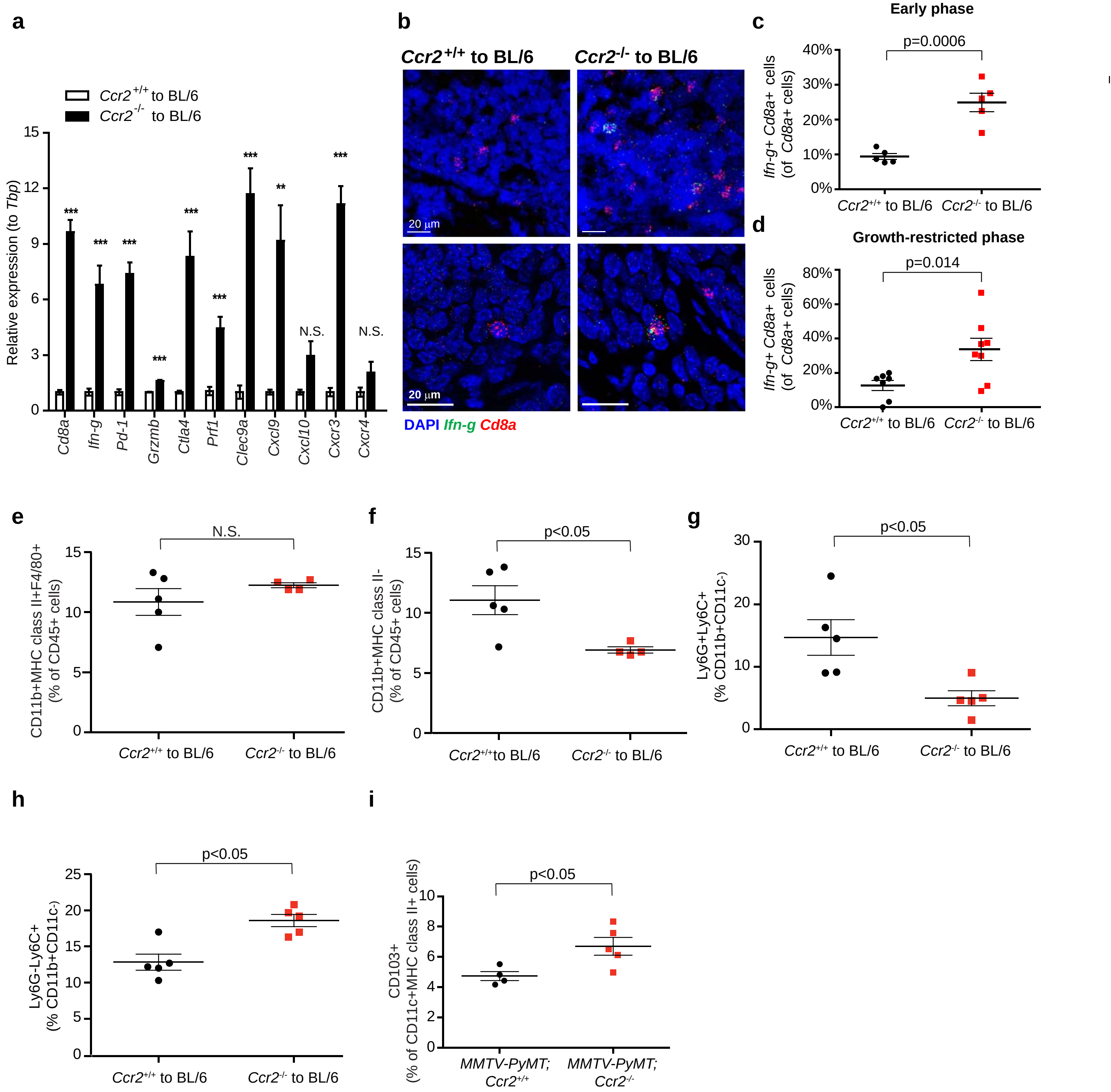

### Supplemental Figure 5

**a**

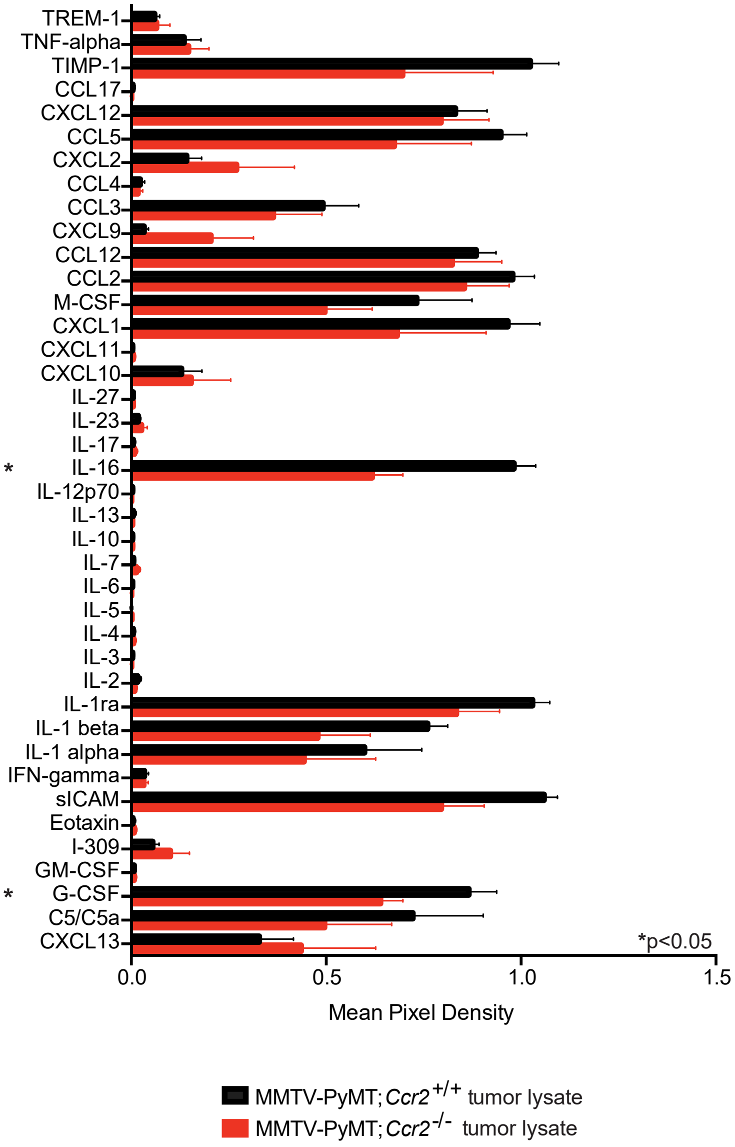


**b**

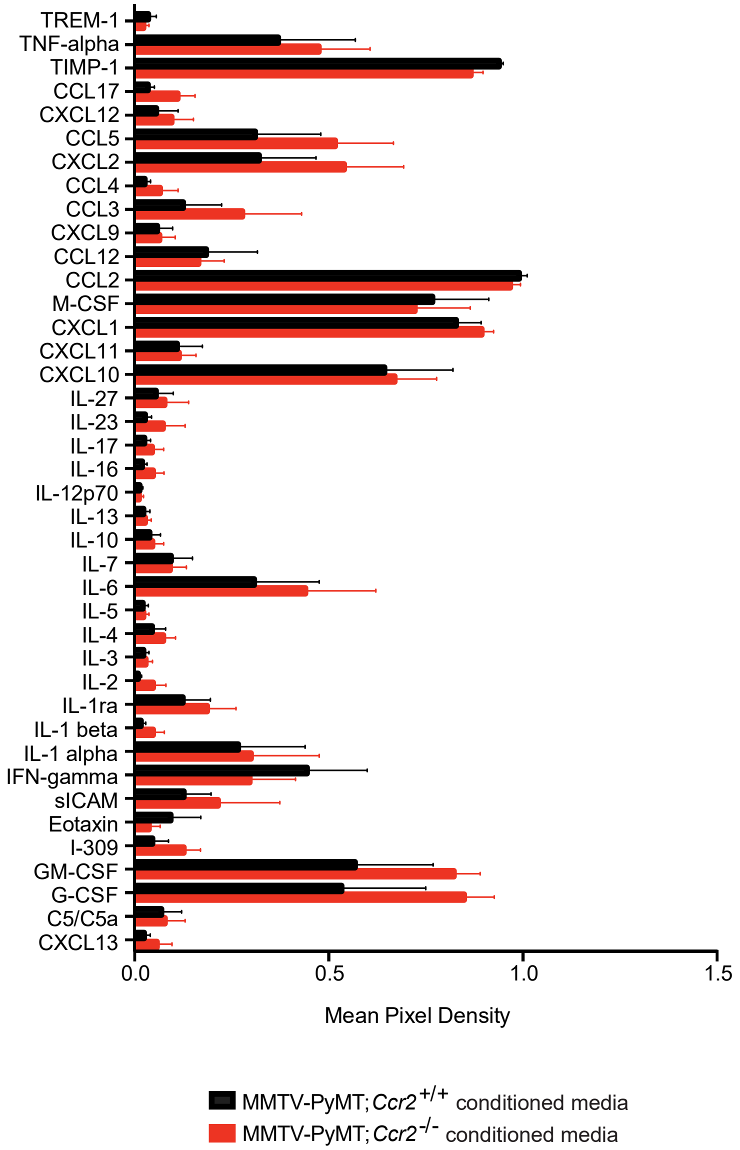
